# Talin mechanosensitivity is modulated by a direct interaction with cyclin-dependent kinase-1

**DOI:** 10.1101/2021.03.19.436208

**Authors:** Rosemarie E. Gough, Matthew C. Jones, Thomas Zacharchenko, Shimin Le, Miao Yu, Guillaume Jacquemet, Ste P. Muench, Jie Yan, Jonathan D. Humphries, Claus Jørgensen, Martin J. Humphries, Benjamin T. Goult

## Abstract

Talin is a mechanosensitive component of adhesion complexes that directly couples integrins to the actin cytoskeleton. In response to force, talin undergoes switch-like behaviour of its multiple rod domains that modulate interactions with its binding partners. Cyclin-dependent kinase-1 (CDK1) is a key regulator of the cell cycle, exerting its effects through synchronised phosphorylation of a large number of protein targets. CDK1 activity also maintains adhesion during interphase, and its inhibition is a prerequisite for the tightly choreographed changes in cell shape and adhesiveness that are required for successful completion of mitosis. Using a combination of biochemical, structural and cell biological approaches, we demonstrate a direct interaction between talin and CDK1 that occurs at sites of integrin-mediated adhesion. Mutagenesis demonstrated that CDK1 contains a functional talin-binding LD motif, and the binding site within talin was pinpointed to helical bundle R8 through the use of recombinant fragments. Talin also contains a consensus CDK1 phosphorylation motif centred on S1589; a site that was phosphorylated by CDK1 *in vitro*. A phosphomimetic mutant of this site within talin lowered the binding affinity of KANK and weakened the mechanical response of the region, potentially altering downstream mechanotransduction pathways. The direct binding of the master cell cycle regulator, CDK1, to the primary integrin effector, talin, therefore provides a primordial solution for coupling the cell proliferation and cell adhesion machineries, and thereby enables microenvironmental control of cell division in multicellular organisms.

**Summary:** The direct binding of the master cell cycle regulator, CDK1, to the primary integrin effector, talin, provides a primordial solution for coupling the cell proliferation and cell adhesion machineries, and thereby enables microenvironmental control of cell division.

## Introduction

Cell adhesion to the extracellular matrix (ECM) is required for anchorage-dependent cell survival and growth in multicellular organisms. During the G1 commitment phase of the cell cycle, adhesion signalling is required to initiate DNA synthesis (1, 2) and suppress apoptosis (3, 4). However, how changes in adhesion signalling are able to influence cell cycle progression in adherent cells is only partly elucidated. During mitosis, major changes in integrin adhesion complexes (IACs), cytoskeletal architecture and cell shape are obligatory for chromosome segregation and cytokinesis (5–8). These remodelling events can be so extensive that cells become round and virtually lose their adhesion. Despite the risks to tissue integrity, the optimally symmetrical geometry of a sphere appears to enable the high degree of precision required for chromosome capture and division plane orientation (9–11). Suppression of these changes during mitosis perturbs the ability of cells to divide accurately (12–14). All of these changes are highly conserved, implying the existence of a primordial regulatory mechanism linking the cell cycle and adhesion machineries.

One link between the cell cycle and adhesion is mediated through the master regulator of the cell cycle, cyclin-dependent kinase-1 (CDK1). CDK1 is a serine/threonine kinase that partners with cyclins that control both kinase activity and substrate specificity (15). CDK1 inhibition leads to disassembly of IACs, suggesting that CDK1 activity has an interphase role in promoting integrin-mediated adhesion and actomyosin organisation (12, 16). IAC area increases during S phase in a CDK1-dependent manner, then inhibition of CDK1 through Wee1-mediated phosphorylation in G2 causes IAC loss in preparation for entry into mitosis (12). Together, these findings identify a key role for CDK1 in the regulation of adhesion during cell cycle progression.

Talins are large (270 kDa) multidomain proteins composed of an N-terminal head coupled to a large rod comprising 13 helical bundles (R1-R13) (17). The two talin isoforms (talin-1 and talin-2) are considered to be the principal proteins that couple integrins to F-actin. The N-terminal FERM domain binds to integrin cytoplasmic domains (18–21) and two sites in the C-terminal flexible rod domain bind F-actin (22, 23). Talin binding to integrins is maintained by actomyosin-generated force and results in conformational activation of integrins (20). The linkage therefore provides a regulatable means of controlling cell adhesion to the ECM from within the cell. This trimolecular core also initiates the recruitment of a large number of additional proteins, termed the adhesome, which varies in composition between IAC type (24–26), reflecting distinct effector, signalling, and mechanosensing functions. The role of talin as a key mechanotransducer relies on its ability to undergo force-dependent structural rearrangements. As force from actin is exerted on talin anchored to integrins, the talin rod gradually unfolds at discrete sites, resulting in both the displacement and recruitment of signalling proteins. This enables talin to serve as a mechanosensitive signalling hub, integrating a wide range of signals to produce diverse cellular responses (27).

Here, we identify a direct interaction between CDK1 and talin through an unbiased analysis of talin binding proteins. CDK1 contains a talin-binding LD motif that interacts directly with the R8 domain of talin. This interaction is required for regulating CDK1 function at IACs and leads to an alteration in the mechanosensitivity of talin. Direct mechanical coupling of the master cell cycle regulator and the principal integrin effector provides an elegant mechanism coupling the cell division and adhesion machineries to facilitate proliferation in a multicellular environment.

## Results

### Talin associates with CDK1 at adhesion sites

To identify novel talin binding proteins, full-length talin-1 fused to GFP was expressed in U2OS cells and interacting proteins identified by GFP-Trap pull-down and mass spectrometry. Multiple proteins were identified, including previously characterised ligands such as vinculin, KANK proteins and integrins (Fig 1A and Supp Table 1). In addition, peptides covering a large proportion of CDK1 (SAINT Score 0.96) were detected with high confidence in all three talin pulldowns, but never in the GFP controls, suggesting that CDK1 also associates with talin. The mass spectrometry data were confirmed by western blotting of the GFP-talin pulldown with anti-CDK1 antibody (Fig 1B). To support these findings, CDK1 was tagged with BirA*, expressed in U2OS cells, and BioID proximity biotinylation employed to label proteins close to CDK1 *in situ*. Using this approach, 118 labelled proteins were identified, including the known CDK1 interactors, cyclin B1 and cyclin B2 (Supp Table 2). 47% of the identified proteins were found within the meta adhesome, suggestive of a role for CDK1 in regulating adhesion/cytoskeletal dynamics (28). Furthermore, in addition to talin, the core adhesome components, vinculin, paxillin, filamin A and CRK, were in close proximity to CDK1 (Fig 1C). Finally, TIRF microscopy was employed to determine the localisation of CDK1 at the cell-ECM interface. mScarlet-CDK1 was distributed in elongated, fleck-like structures that partially overlapped with GFP-talin-containing IACs (Fig 1D). Taken together, these findings indicate a close association between talin and CDK1 at sites of integrin-mediated adhesion to the ECM.

**Figure 1:**
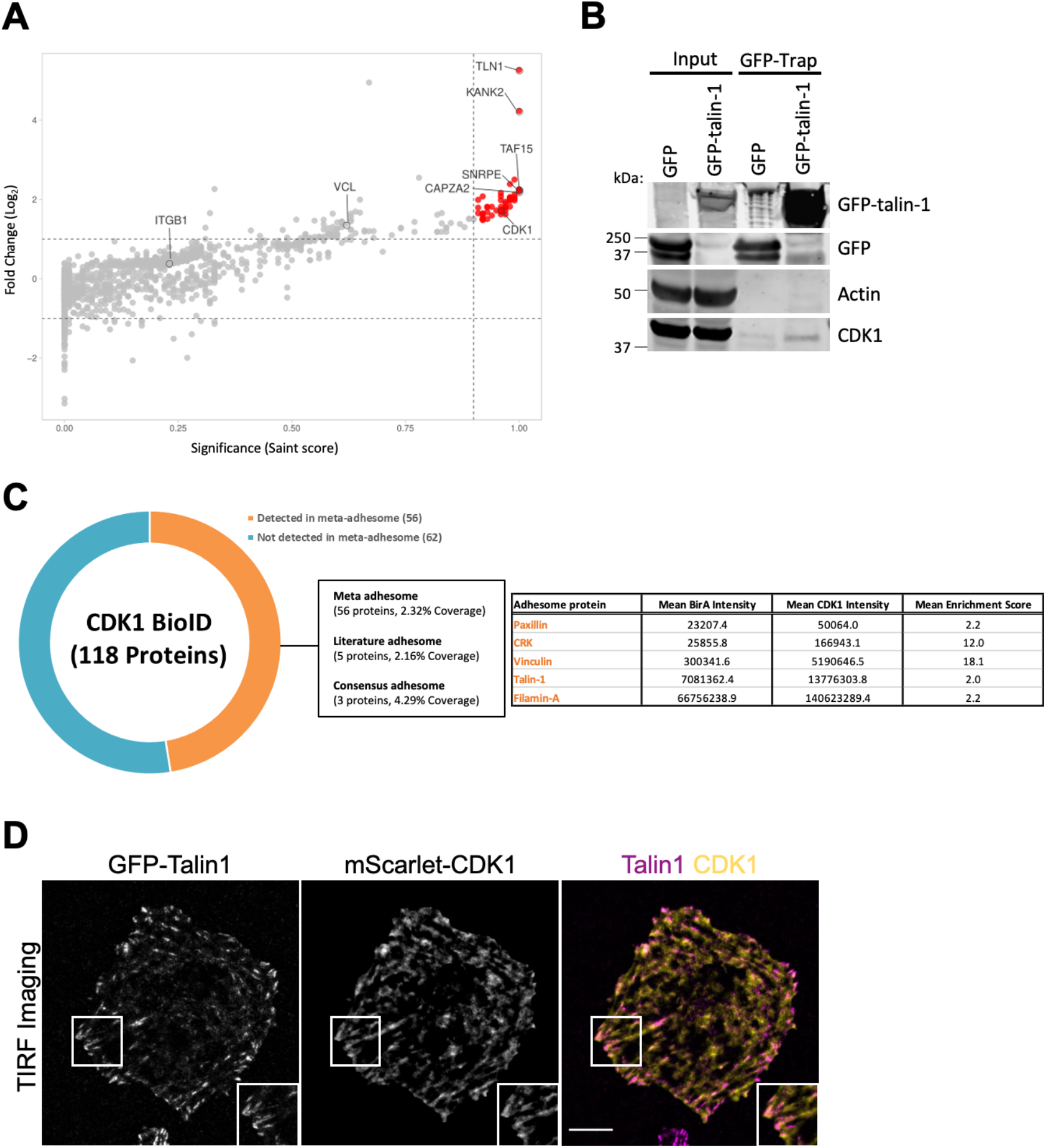
Talin interacts with CDK1. (**A**) MS analysis of GFP-tagged talin-1 binding proteins, displayed as a volcano plot. The fold-change enrichment over GFP alone is plotted against the significance of the association. The SAINT score and the FC_B scores were calculated using the REPRINT (Resource for Evaluation of Protein Interaction Network) online tool (https://reprint-apms.org/). The volcano plot was generated using VolcaNoseR (29). Key putative talin-1 binders, including CDK1, are highlighted. (**B**) GFP-Trap pulldown of GFP alone or GFP-talin-1 expressed in U2OS cells followed by western blotting for GFP, actin and CDK1. (**C**) MS analysis of proteins biotinylated by BirA-CDK1, indicating proteins in close proximity to CDK1 in cells. Proteins in table are core components of IACs. (**D**) TIRF image of a U2OS cell co-expressing GFP-talin-1 and mScarlet-CDK1. Scale bar 10 μm.

### CDK1 contains an LD motif that binds directly to talin R8

We next aimed to identify the binding interface between talin and CDK1. Talin contains multiple binding sites for proteins that contain leucine/aspartic acid LD motifs, with a consensus sequence I/LDxØØxØØ (where Ø denotes a hydrophobic residue) (30, 31). These amphipathic helical peptide motifs engage talin via a helix-addition mechanism (32, 33), packing on the side of the helical bundles in the talin rod. This mode of binding allows mechanosensitive regulation as it only occurs when the talin rod domain mediating the interaction is folded: unfolding due to mechanical force disrupts the LD motif binding site and the connection is severed (27).

Analysis of the CDK1 protein sequence identified a highly conserved, consensus LD motif sequence between residues 206-223 (Fig 2A,B), which we postulated might bind directly to talin. Fluorescence polarisation was employed to test direct binding, as described previously (34, 35). In this assay, a synthetic peptide spanning the CDK1 LD motif, CDK1(206-223), is fluorescently-tagged and titrated against an increasing concentration of unlabelled talin fragment; any binding between the two polypeptides results in an increase in the fluorescence polarisation signal. To identify the putative CDK1 binding site(s) on talin, six talin fragments were generated that span the whole molecule; F0-F3, R1-R3, R4-R8, R9-R10, R11-R12 and R13-DD (Fig 2C). Two of these fragments incorporate the actin-binding sites in the talin rod, ABS2 (R4-R8 (23, 36)) and ABS3 (R13-DD (37, 38)). No significant interaction was observed between CDK1(206-223) and the talin Head, R1-R3, R9-R10, R11-R12 or R13-DD fragments; however, R4-R8 demonstrated increasing fluorescence polarisation with increasing talin concentration, indicative of a direct interaction (Fig 2D). To further define the CDK1 binding site within talin R4-R8, we tested binding to additional rod domain constructs. Talin R4-R6 demonstrated negligible binding to CDK1(206-223), whereas R7R8 bound to CDK1(206-223) with a Kd of 14 μM (Fig 2E). The R7R8 region of both talin isoforms, talin-1 and talin-2, bound CDK1(206-223) with similar affinity (Fig 2E). As found for other LD motif peptides (35), mutation of the LD motif within CDK1 to two alanine residues (to generate a 2A mutant; Fig 3A,B) markedly reduced the interaction with talin (Fig 3C).

**Figure 2:**
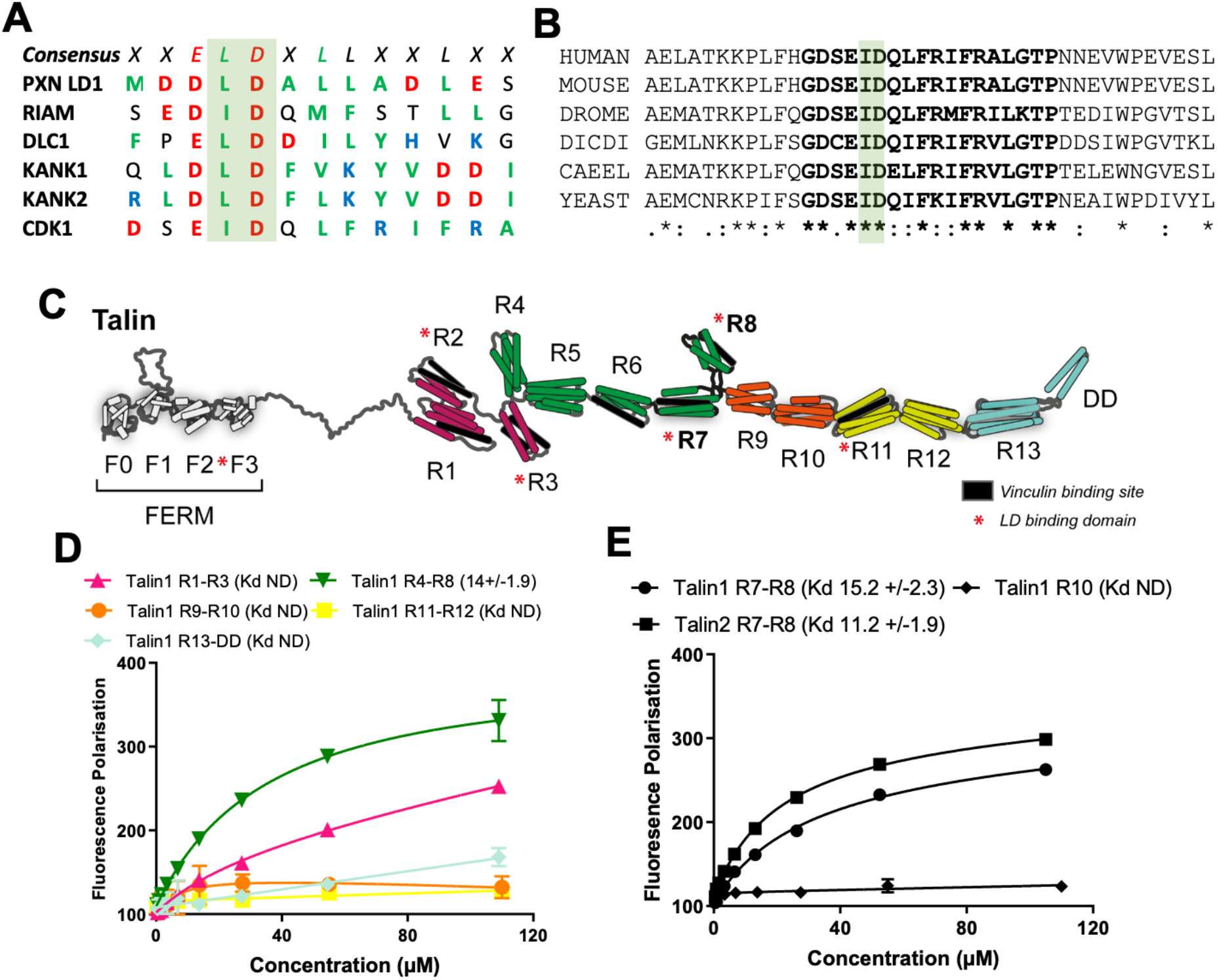
CDK1 interacts with the talin R8 domain. (**A**) Sequence alignment of the previously reported talin-binding LD motifs, paxillin LD1, RIAM, DLC1, KANK1 and KANK2, with CDK1. Acidic (red), basic (blue) and hydrophobic (green) residues are highlighted. (**B**) Sequence alignment of the CDK1 region containing the talin binding site. The talin-binding LD motif is highlighted in bold. Aligned sequences were human (UniProt P06493), mouse (UniProt P11440), *Drosophila melanogaster* (UniProt P23572), *Dictyostelium discoideum* (UniProt P34112), *Caenorhabditis elegans* (UniProt P34556) and *Saccharomyces cerevisiae* (UniProt P00546). (**C**) Schematic of the domain structure of talin. The five talin rod fragments screened for binding to CDK1 are highlighted: R1-R3 (residues 482-911, pink), R4-R8 (residues 913-1653, green), R9-R10 (residues 1655-2015, orange), R11-R12 (1974-2293, yellow) and R13-DD (2300-2542, cyan). The 11 vinculin binding sites are shown in black. The LD binding domains identified to date are shown by a red asterisk. (**D-E**) Binding of BODIPY-labelled CDK1(206-223) peptide to talin fragments measured using fluorescence polarisation. Binding of BODIPY-labelled CDK1(206-223) peptide with (**D**) the five talin fragments and (**E**) with talin-1 and talin-2 R7R8 domains. Dissociation constants ± SE (μM) for the interactions are indicated in the legend. All measurements were performed in triplicate. ND = not determined.

**Figure 3:**
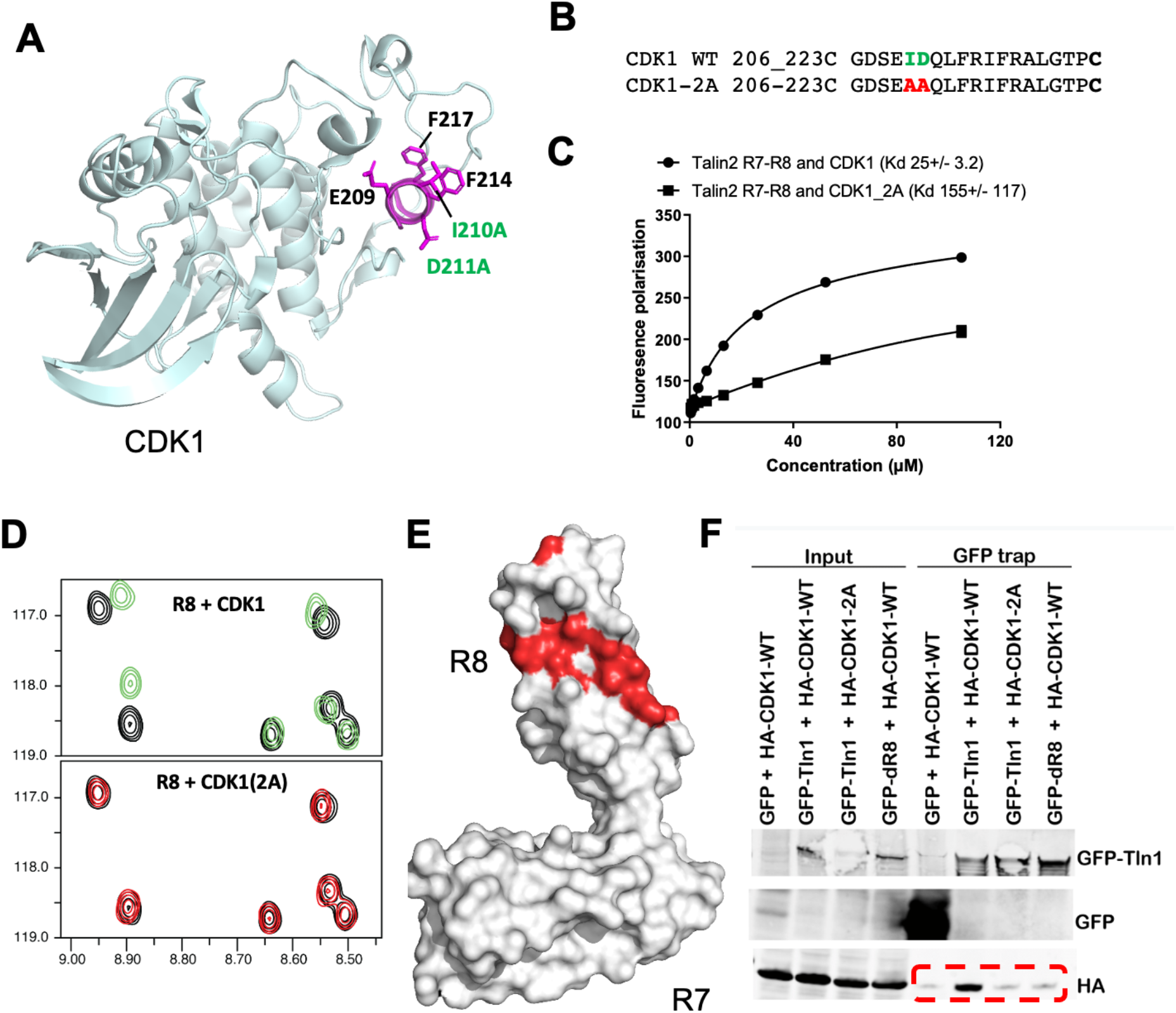
The CDK1 LD motif is required for CDK1 binding to talin-1. (**A**) CDK1 structure (PDB ID: 4YC3 (39)) with the talin-binding LD motif highlighted in magenta. The LD motif consensus residues (E209, I210, D211, F214 and F217) and the I210A and D211A, 2A mutation are highlighted. (**B**) Aligned sequences of CDK1 WT and CDK1-2A mutant peptides. (**C**) Binding of BODIPY-labelled CDK1 WT and 2A 206-223 peptide with talin-1 R7R8 domains using fluorescence polarisation. Dissociation constants ± SE (μM) for the interactions are indicated. All measurements were performed in triplicate. (**D**) ^1^H,^15^N HSQC spectra of 70 μM ^15^N-labelled talin-1 R8 (residues 1461-1580) (black) with 420 μM CDK1 peptide (green), 420 μM CDK1-2A peptide (red) added. (**E**) Mapping of the CDK1 binding site on R8 as detected by NMR using weighted chemical shift differences (red) – mapped onto the R7R8 structure. (**F**) GFP-Trap pulldowns from cells expressing GFP alone, GFP-WT-talin-1 or GFP-ΔR8-talin-1 along with HA-tagged WT or 2A-CDK1, followed by western blotting for GFP and HA.

### The CDK1 LD motif binding pocket on talin R8 differs from that of DLC1 and RIAM

We initially used NMR to characterise the interaction between CDK1 and talin. HSQC spectra of ^15^N-labeled talin-1 R7R8 were collected in the absence and presence of increasing amounts of CDK1(206-223) peptide (Fig 3D). Addition of CDK1(206-223) peptide, but not CDK1-2A, resulted in progressive chemical shift changes to a subset of the R8 residues, indicative of a direct, specific interaction. Using the chemical shift assignments of the R8 domain of talin-1 (BMRB ID:19339 (33)), the changes were mapped onto the structure of R8 (PDB ID: 2X0C (40)). The changes mapped onto one face of the R8 domain, the same region that binds the LD motifs of DLC1 (33) and RIAM (17, 41) (Fig 3E). To support the requirement for the talin R8 domain in interacting with CDK1, GFP-Trap pulldowns were performed using wild-type (WT) full-length talin or talin lacking the R8 domain (ΔR8). WT-talin-1-GFP pulled down HA-tagged CDK1-WT, but not HA-CDK1-2A, whereas ΔR8-talin-1-GFP failed to associate with HA-CDK1-WT (Fig 3F).

To resolve the structural basis of the talin-CDK1 interaction, the R7R8-CDK1 peptide complex was crystallised and the structure determined to 2.28Å resolution (Fig 4A,B). The structure contained two R7R8 molecules within the asymmetric unit, each with CDK1 peptides localised to the respective R8 domains (Supp Fig 1A; Supp Table 3). Each peptide was exceptionally resolved in both the F_0_-F_C_ weighted difference map and simulated annealing OMIT maps (Supp Fig 1B,C). Unexpectedly, the CDK1 peptide was oriented perpendicular to the helices of the R8 fold, forming a pseudo-5-helix bundle (Fig 4A,B) that contrasts with the helix-addition mechanism of DLC1 and RIAM (33, 41) (Fig 4 C,D). This arrangement was supported by NMR chemical shift mapping (Fig 3E), where the shift changes upon addition of CDK1 peptide were located predominantly at one end of the LD motif-binding surface of the bundle.

**Figure 4:**
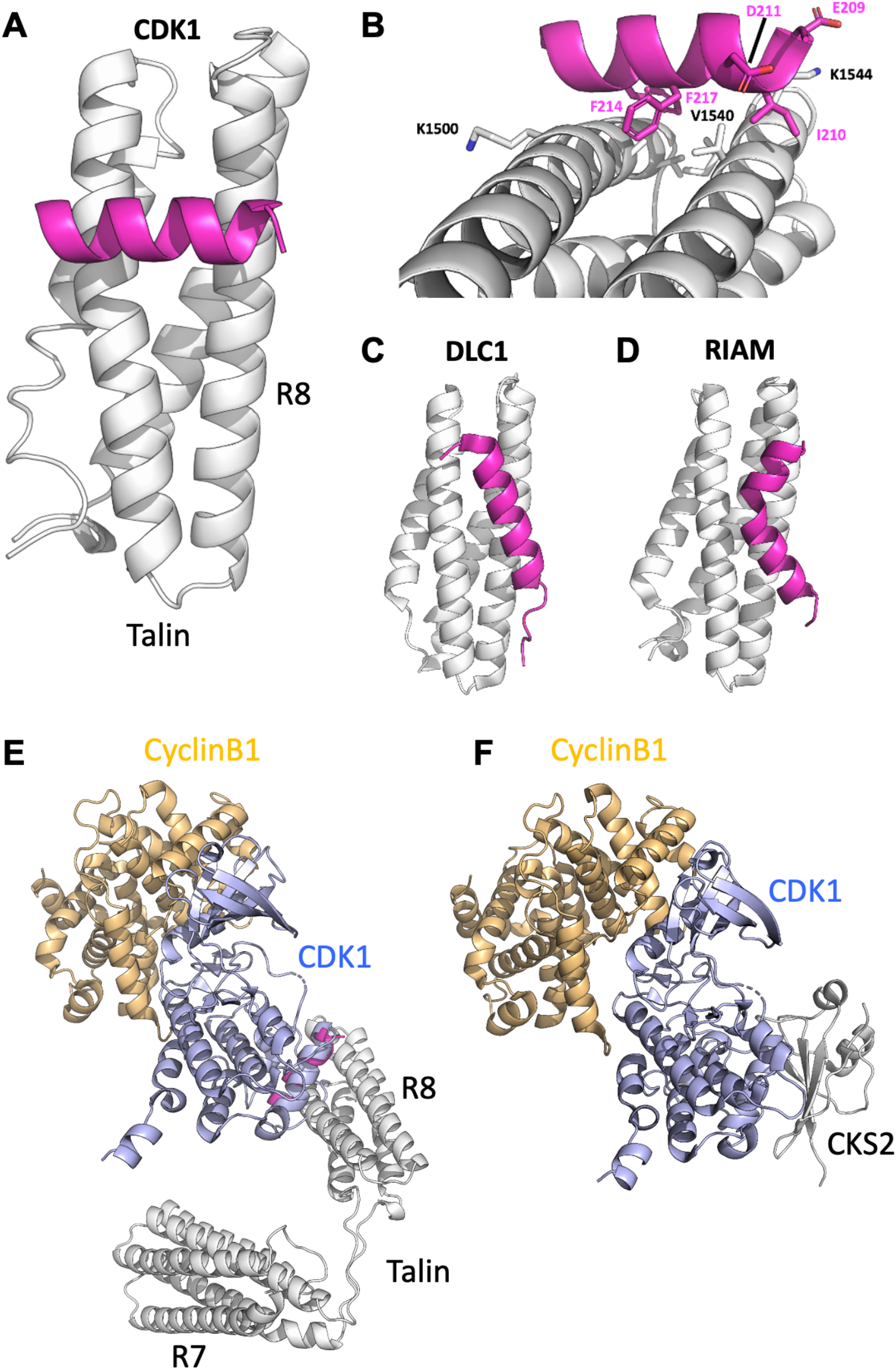
Crystal structure of the talin-CDK1 complex. (**A-B**) Cartoon representation of the X-ray structure of the talin R7R8 fragment complex with CDK1(206-223). Talin R8 (white) and CDK1 (magenta) are indicated. (A) Face on, and (**B**) side on views. Key residues are highlighted. (**C-D**) Structure of talin R8-LD motif complexes. (**C**) Talin-RIAM (PDB ID: 4W8P (41)) and (**D**) talin-DLC1 (PDB ID: 5FZT (33)), orientated as in (**A**). (**E-F**) Structural model of a talin-CDK1-cyclin complex.(**E**) Modelled structure of talin (grey) bound to the complex of CDK1 (blue) and cyclin B1 (orange) using (PDB ID: 4YC3 (39)). (**F**) Structure of the tripartite complex of CDK1 (blue), cyclin B1 (orange) and CKS2 (grey) (PDB ID: 4YC3). Talin R8 and CKS2 both bind CDK1 via the same region on the opposite face to the cyclin binding surface.

The CDK1 peptide was identified on the basis of sequence homology with the LD motif. In previous studies of LD motif recognition, the aspartate residue of the LD motif was shown to form a salt bridge with a positively charged residue on the cognate binding site (42) and, in the case of DLC1, mutation of talin K1544 abolished interactions with R8 (33). The CDK1-talin structure deviated from this shared interaction mechanism as K1544 was not involved in the interaction, but instead oriented away from the CDK1 binding site. The isoleucine side chain of the LD motif was responsible for displacing the K1544 side chain and an ID/AA double mutation attenuated the CDK1-talin interaction (Fig 4B). The remainder of the CDK1 peptide formed a hydrophobic interface with the R8 domain with the CDK1 ^214^FRIFRA^219^ sequence buried into a hydrophobic groove between the helices formed of talin residues L1492, A1495, V1498, L1539, V1540 and I1543 (Fig 4B).

The CDK1 sequence formed an amphipathic helix that closely resembled its conformation in full-length CDK1 (PDB ID: 4YC6 (39)), where the key residues involved in talin binding are surface exposed (Supp Fig 2). The structure of this lobe of the kinase has been solved in complex with other proteins such as CKS1 (cyclin-dependent regulatory subunit 1) (39), suggesting it may play a role as a targeting domain. As the CDK1 ID-motif peptide superimposed on the CDK1 structure, it was possible to model the interaction between talin and the CDK1-cyclin B1 complex (Fig 4E,F). This demonstrated that the talin-CDK1 interaction is unlikely to perturb CDK1 binding to regulatory cyclin proteins and suggests the possibility that talin is able to interact with active CDK1-cyclin complexes.

### Talin binding is required for CDK1-dependent regulation of adhesion

We previously demonstrated that CDK1 kinase activity is required to maintain IACs and to facilitate changes in IAC area during cell cycle progression (12). To test whether these functions were dependent on the interaction between CDK1 and talin, the effects of mutant versions of both proteins were determined. Consistent with previous data, knockdown of CDK1 in asynchronous cells resulted in a reduction in paxillin-positive IAC area that was rescued by re-expression of siRNA-resistant WT CDK1, but not CDK1-2A (which is unable to bind talin; Supp Fig 3A,B). In addition, in synchronised cells expressing CDK1-WT, the robust IAC growth previously observed during S phase did not occur in cells expressing CDK1-2A (Supp Fig 3C). These findings indicate that the CDK1 LD motif is required for CDK1 to maintain IACs in asynchronous cells and to facilitate IAC growth during S phase. Subsequent analysis of the CDK1-2A mutant protein demonstrated complexing with cyclin B1, but not cyclin A2, and a lack of phosphorylation at the activating site T161 (Supp Fig 3D,E). While talin binding may therefore contribute to CDK1 activation, since CDK1-cyclin A2 maintains IACs (9), it therefore cannot be concluded that the effects of CDK1-2A on adhesion are solely attributable to the loss of CDK1 binding to talin.

In complementary studies, the effects of ΔR8-talin-1 expression on IACs were examined. In cells expressing WT talin-1, a dose-dependent reduction in IAC area was observed following treatment of cells with the CDK1 inhibitor RO3306. In contrast, no loss of IAC area was observed in cells expressing ΔR8-talin-1 (Fig 5A,B). Furthermore, in synchronised cells expressing ΔR8-talin-1, IAC area remained constant through G1, S and G2, while cells expressing WT talin-1 displayed IAC growth during S (Fig 5C). These data demonstrate that the talin R8 domain is required for facilitating CDK1-dependent regulation of IACs.

**Figure 5:**
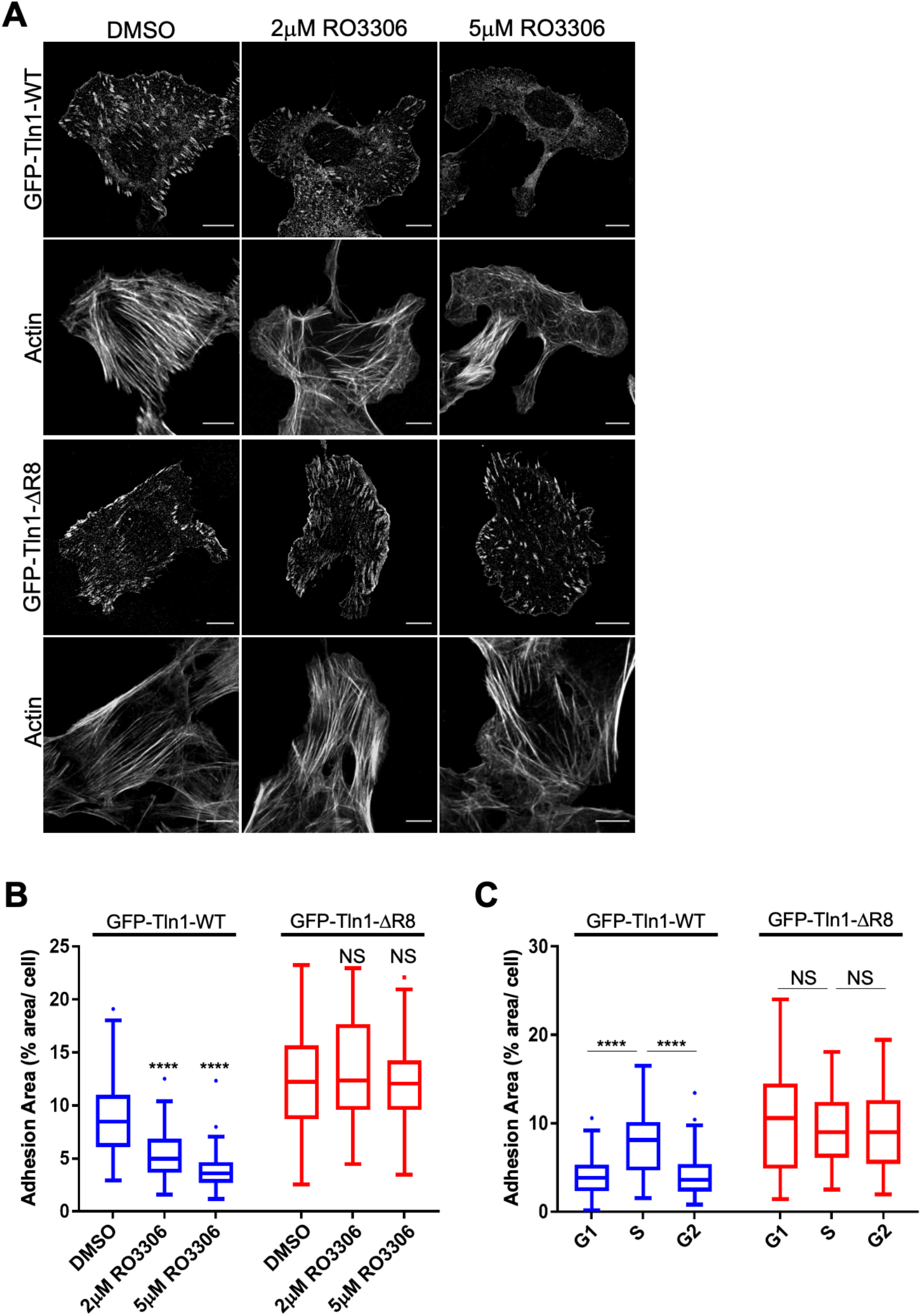
Talin-CDK1 interaction is required to facilitate CDK1-dependent regulation of adhesion complexes. (**A**) Confocal imaging of U2OS cells expressing GFP-WT-talin-1 or GFP-ΔR8-talin-1 treated with DMSO or two different doses of the CDK1 inhibitor RO3306. Bars 10 μm. (**B**) IAC area changes in cells treated with DMSO or RO3306. (**C**) IAC area changes in G1, S, and G2 phase for cells expressing GFP-WT-talin-1 or GFP-ΔR8-talin-1. For B and C, a minimum of 50 cells per condition was used for analysis and results are displayed as Tukey box and whisker plots (whiskers represent 1.5x interquartile range). ****, p<0.0001.

### CDK1 phosphorylates talin R7R8 and modulates its mechanosensitivity

Talin R8 binds to CDK1 in a similar region to the regulatory CKS proteins that target CDK1 to specific substrates and facilitate CDK1-dependent phosphorylation (39, 43). We therefore hypothesised that binding of CDK1 to talin might lead to phosphorylation of residues within the R7R8 region. Incubation of talin-1 R7R8 and CDK1-cyclin A with ATP *in vitro* resulted in a high level of phosphorylation of R7R8 (Fig 6A,B), and subsequent analysis by mass spectrometry identified a single phosphorylation at S1589. Furthermore, sequence analysis identified the presence of a consensus CDK1 phosphorylation motif (SP) at S1589 (Fig 6B-D).

**Figure 6:**
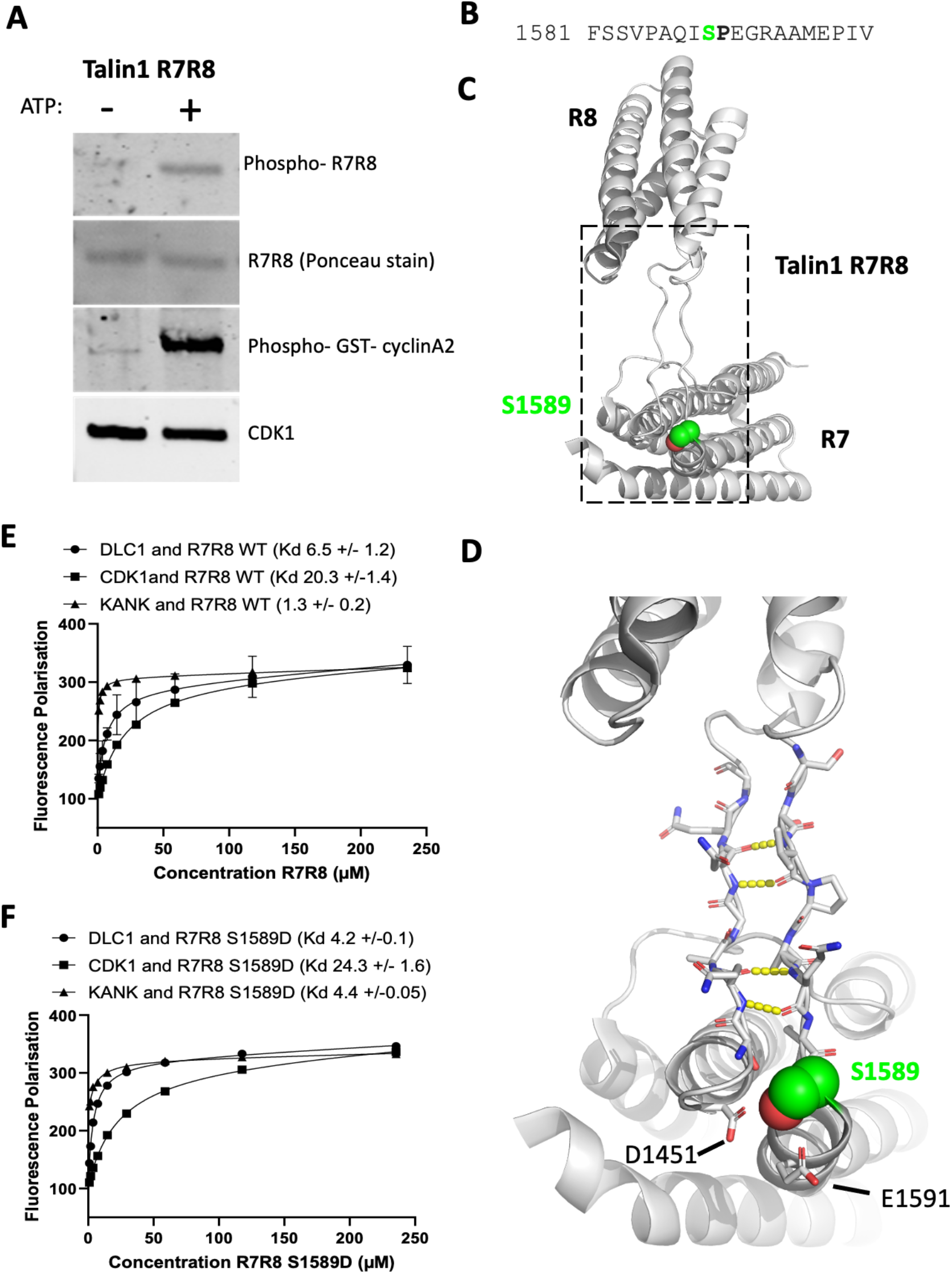
CDK1-cyclin A2 phosphorylates talin-1. **(A)** Western blotting of talin-1 R7R8 phosphorylation by purified CDK1-cyclin A2 in the presence of ATP. **(B)** Talin-1 sequence highlighting the SP motif that contains the phosphorylation site at S1589. **(C)** Talin-1 R7R8 structure with the phosphorylation site, S1589, highlighted in green. (**D)** The linker between the R7 and R8 domains forms a two-stranded anti-parallel ß-sheet-like structure mediated by a hydrogen bonding network (dashed yellow lines). The location of the S1589 residue (green spheres) relative to the acidic side chains of D1451 and E1591. (**E-F**) Binding of BODIPY-TMR labelled CDK1 206-223C, DLC1 465-489C and KANK1 30-60C peptides to talin-1 R7R8 (1357–1653) (**DE**) WT and (**F**) S1589D. Binding affinities were measured by fluorescence polarization. Dissociation constants ± SE (μM) for the interactions are indicated.

The positioning of the talin-1 phosphorylation site is at the end of the R7 rod domain close to where R8 is inserted (Fig. 6C-D), and is adjacent to two charged residues D1451 and E1591. We predicted that phosphorylation of S1589 might lead to repulsion from these acidic residues, resulting in decreased stability of both R7 and R8 and a potential weakening of the binding of ligands to R7R8. To determine if S1589 phosphorylation affected the binding of KANK to R7 and DLC1 and CDK1 binding to R8 (33, 35), a talin phosphomimetic mutant (S1589D) was designed and the binding affinity of the CDK1, KANK and DLC1 peptide ligands to R7R8 measured by fluorescence polarisation. Binding of both R8 ligands was largely unaffected by the talin-1 S1589D mutation, but the affinity of KANK binding was approximately four-fold lower (Kd 4 μM and 1 μM for the mutant and WT, respectively; Fig 6E,F). These data indicate that S1589 phosphorylation does not appreciably alter the binding surface of R8, but does have a potentially significant change in binding of KANK to R7.

The stability and positioning of the R7 and R8 domains is maintained by an extensive network of hydrogen bonds between the linkers connecting the two domains forming a two-stranded, anti-parallel ß-sheet-like structure (40). As S1589 is located close to the linker between R7 and R8, it was conceivable that the mechanosensitivity of this region may be altered by CDK1-dependent phosphorylation. To test whether post-translational modification of S1589 alters the mechanical stability of the domains, we used single molecule unfolding experiments using magnetic tweezers (Fig. 7A). Our previous study of the mechanical response of talin showed that, when exposed to mechanical force, the R7R8 double domain unfolds cooperatively with a single unfolding step of ~80 nm at ~15 pN force at a force loading rate of ~3.4 pN s^−1^ (44). To further characterise the mechanical properties of R7R8, single molecule unfolding experiments were conducted for an R7R8 construct at loading rates of 1.0±0.1 and 5.0±.5 pN s^−1^. Consistent with our previous report (44), a cooperative unfolding event was observed indicated by a single height-jump step of ~70 nm at a force of ~14 pN (Fig 7B). The normalised unfolding force distributions of R7R8 at force loading rates of 1±0.1 pN s^−1^ and 5±0.5 pN s^−1^ peaked at 13.3±1.6 pN and 14.5±1.9 pN, respectively. Next, similar force-increase scans were performed with a single molecule construct containing R7 alone (Fig 7C). In this case, the unfolding forces of the R7 domain were 10.1±1.9 pN and 11.2±1.1 pN for 1±0.1 pN s^−1^ and 5±0.5 pN s^−1^, respectively (Fig 7C), demonstrating that the R7 domain is mechanically weaker than R7R8. As it has already been demonstrated that R8 is mechanically weak (44, 45), these findings indicate that the interdomain interaction between R7 and R8 is mutually stabilising.

**Figure 7:**
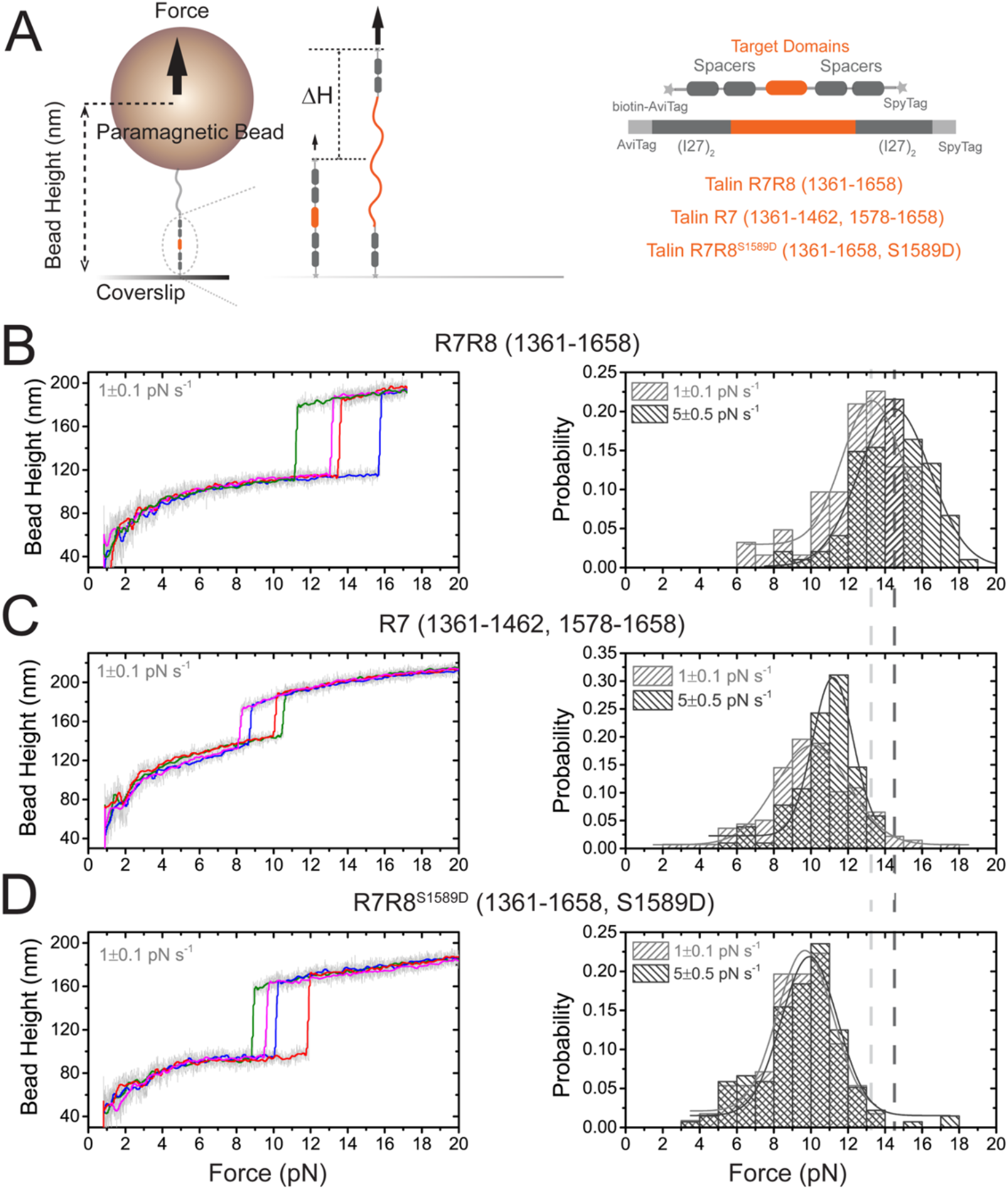
R7R8 module is mechanically stabilised by R7-R8 interdomain interaction. (**A**) Schematic of single molecule stretching experiments where a single protein tether, coupled between a paramagnetic bead and a surface, is stretched under force. (**B**) **Left panel**: Typical force-bead height curves of the R7R8 construct during force-increase scans at a loading rate of 1 pN s^−1^. Each coloured line indicates one independent force-increase scan, smoothed (10-point fast Fourier transform) from raw data (grey). **Right panel**: The normalised unfolding force distributions of R7R8 complex during force-increase scans at 1 pN s^−1^ and 5 pN s^−1^. The data points obtained for analysis are N=62 and N=195, respectively. The curves are Gaussian fitting of the distribution with peaks at 13.3±1.6 pN and 14.5±1.9 pN, respectively. (**C**) **Left panel**: Typical force–bead height curves of the R7 with short linker region during force-increase scans at a loading rate of 1 pN s^−1^. **Right panel**: The normalised unfolding force distributions of R7 domain during force-increase scans at 1 pN s^−1^ and 5 pN s^−1^. The data points obtained for analysis are N=57 (1 pN s^−1^) and N=65 (5 pN s^−1^), respectively. The curves are Gaussian fitting of the distribution with peaks at 7.3±1.1 pN and 9.5±1.1 pN, respectively. (**D**) **Left panel**: Typical force–bead height curves of R7 with a long linker during force-increase scans at a loading rate of 1 pN s^−1^. **Right panel**: The normalised unfolding force distributions of R7 domain during force-increase scans at 1 pN s^−1^ and 5 pN s^−1^. The data points obtained for analysis are N=138 (1 pN s^−1^) and N=103 (5 pN s^−1^), respectively. The curves are Gaussian fitting of the distribution with peaks at 10.1±1.9 pN and 11.2±1.1 pN, respectively. (**E**) **Left panel:** Typical force–bead height curves of the R7R8^S1598D^ construct during force-increase scans at a loading rate of 1 pN s^−1^. **Right panel**: The normalised unfolding force distributions of R7R8^S1598D^ complex during force-increase scans at 1 pN s^−1^ and 5 pN s^−1^. The data points obtained for analysis are N=112 (1 pN s^−1^) and N=136 (5 pN s^−1^), respectively. The curves are Gaussian fitting of the distribution with peaks at 9.7±1.5 pN and 10.0 ±1.5 pN, respectively. The light grey and the dark grey dashed lines indicate the corresponding peak force values for R7R8 at 1 pN s^−1^ and 5 pN s^−1^, respectively.

**Figure 8:**
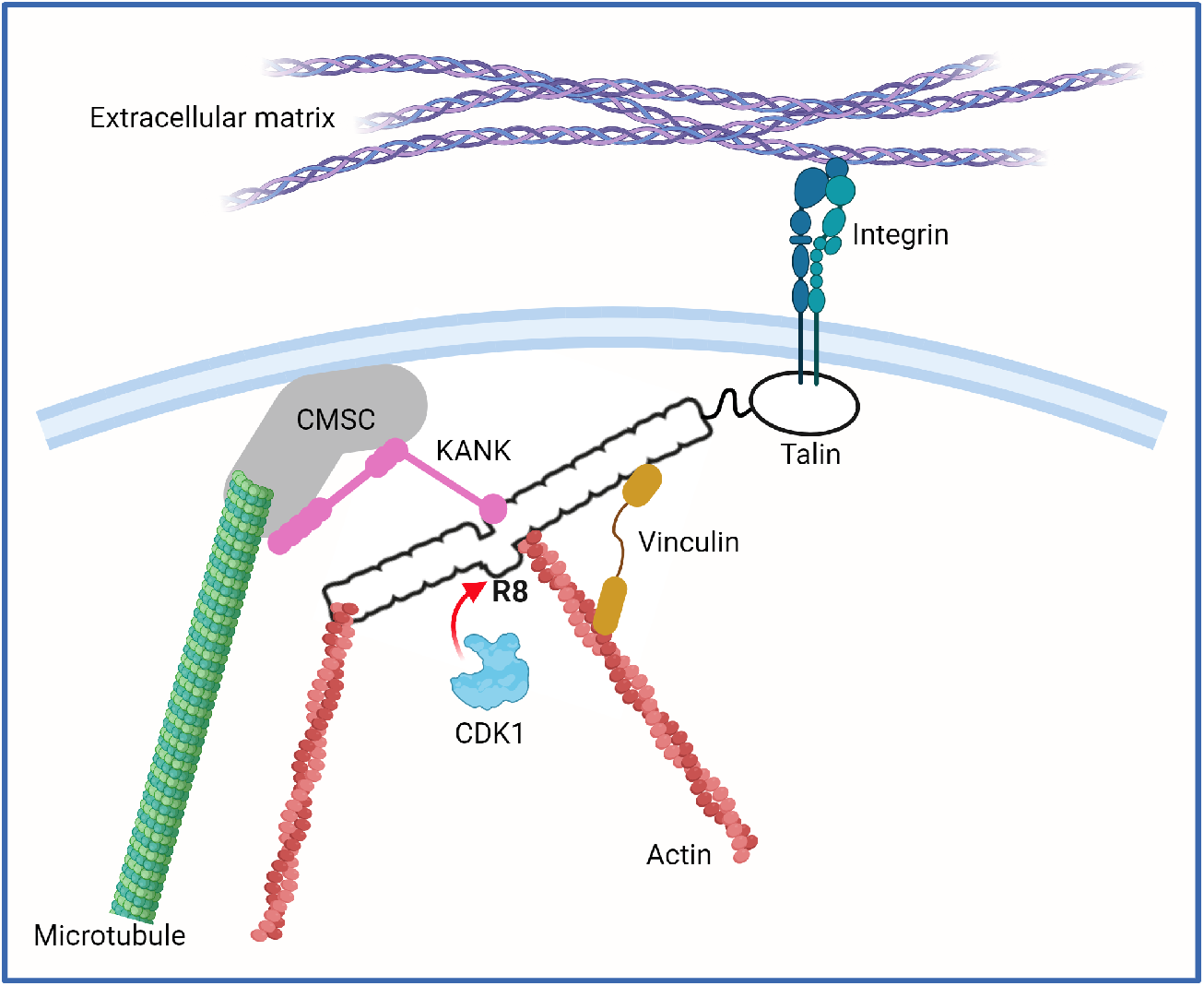
A cartoon to show how CDK1 regulation of talin mechanical response might orchestrate global rearrangements of the cells cytoskeleton. Talin (white) acts as a Mechanosensitive Signalling Hub that coordinates multiple cytoskeletal connections including microtubules (green), mediated via KANK (pink) and the cortical microtubule stabilising complex (CMSC; orange) and actin (red), both directly and indirectly via vinculin (green). The cell signalling of each integrin adhesion complex will be dependent on the conformational patterns of the talin binary switch domains and the cytoskeletal connections. The R7R8 region of talin serves as a nexus between the integrin adhesion complexes, microtubules and actin filaments. CDK1 (blue) binding to, phosphorylating, and altering the mechanical response of the R7R8 domains has the potential to alter these cytoskeletal linkages which might trigger global cytoskeletal rearrangement.

Next, to investigate how phosphorylation might affect the mechanical stability of R7R8, a phosphomimetic single molecule construct of R7R8 containing S1589D (referred as R7R8^S1589D^) was generated. The unfolding force of R7R8^S1589D^ peaked at 9.7±1.5 pN and 10.0 ± 1.5 pN for 1±0.1 pN s^−1^ and 5±0.5 pN s^−1^, respectively, and was significantly reduced compared to WT R7R8 (Fig 7D). Furthermore, R7R8^S1589D^ and R7 had similar mechanical stabilities. These results demonstrate that phosphorylation of S1589 weakens the mechanical stability of this region likely by disrupting the interdomain connection of R7 and R8.

## Discussion

Much of our current understanding of cell cycle regulation stems from studies on unicellular organisms, and less is known about the mechanisms coordinating cell division in the extracellular matrix-containing tissues of multicellular organisms. Here, our principal discovery is that there is a direct binding interaction between the master cell cycle regulator, CDK1, and talin, a large mechanosensitive, actin-binding protein that plays a key role in regulating integrin-mediated cell adhesion to extracellular matrices. We have employed biochemical and structural approaches to define the binding mechanism, and demonstrate that a helical, talin-binding LD motif in CDK1 engages the talin rod R8 helical bundle. A CDK1-2A mutation, which disrupts the LD motif, impedes the assembly of IACs. Talin also contains a functional consensus CDK1 phosphorylation motif centred on S1589, which regulates the mechanical responsiveness of the region, and thereby potentially alters downstream mechanotransduction pathways. We hypothesise that CDK1 binding to the talin scaffold could be a driver of the major morphological changes in adhesion seen during cell cycle progression and division, and that this constitutes an evolutionary adaptation of the cell cycle that is fundamental to multicellularity.

Although we demonstrate that the talin-CDK1 interaction is required for CDK1-dependent regulation of IACs during the cell cycle, a more detailed understanding of the mechanisms underlying this effect remains to be elucidated. For example, the talin-CDK1 interaction might regulate phosphorylation of multiple adhesome components, thereby contributing to IAC dynamics and downstream signalling pathways. The talin R8 domain is a nexus for multiple binding interactions; thus, it binds DLC1 (33, 46), RIAM (17, 47) and paxillin (33), which modulate various aspects of cell spreading and migration, raising the possibility that competing interactions might provide a novel regulatory axis. The unique structure of the R7R8 region, in which the R8 helical bundle is inserted into the R7 helical bundle, means that R8 is shielded from force, unfolding only when R7 unfolds at ~14 pN applied force (44, 45). R8 therefore retains its ability to form helical addition interactions with LD motif-containing proteins even when flanking talin domains have unfolded under the forces exerted at sites of adhesion. The R7R8 region also provides linkages between IACs and the cytoskeleton, and it connects to both the actin and microtubule networks (Fig 7). Specifically, the R7 domain binds the KANK family of proteins (35, 48), which link to the cortical microtubule stabilising complex, and the R8 domain forms part of the actin binding site 2 (ABS2) that makes strong, tension-bearing linkages to the actin cytoskeleton (23, 36). The R7R8 domains are stabilised by a ladder of hydrogen bonds between the linkers adjoining the two domains (40), which makes the double domain one of the more stable regions in the group III (15-21 pN) of talin rod domains (44). The S1589D phosphomimetic R7R8 mutant is more susceptible to force, and unfolds at ~10 pN. This raises the possibility that CDK1 binding to, and phosphorylation of, the R7R8 region might regulate the association of talin with other IAC components and the cytoskeleton. The accessibility of the VBS in R8 has recently been shown to facilitate maturation of nascent adhesions (49), suggesting that phosphorylation of S1589 during the cell cycle might also regulate adhesion dynamics by controlling the availability of this helix.

The talin binding site on CDK1 also binds to the CKS proteins that regulate CDK1 function (39). Therefore, talin and CKS interactions with CDK1 should be mutually exclusive. It is possible that the part of the CDK1 catalytic domain that binds to talin and CKS is a CDK1-targetting domain, and its interactions with various proteins relocate CDK1 to cellular compartments where it can exert its effects. In addition, mutating the CDK1 LD motif leads to a loss of CDK1 phosphorylation at T161, suggesting that this motif plays a key role in upstream regulation of CDK1 function. Therefore, investigating how binding partners of CDK1-WT and CDK1-2A differ could elucidate key CDK1 regulatory mechanisms. Binding of CDK1 to cyclin A requires prior phosphorylation of T161, whereas binding of CDK1 to cyclin B occurs without this phosphorylation (50), and we demonstrate that CDK1-2A is still able to associate effectively with cyclin B, but is unable to complex with cyclin A. Thus, the CDK1-2A mutant may prove to be a useful tool for distinguishing the different roles of CDK1-cyclin A and CDK1-cyclin B during cell cycle progression. In this context, expression of CDK1-2A leads to a reduction in IACs and loss of IAC growth during S phase, processes that we have previously attributed to CDK1-cyclin A, implying its utility in elucidating the mechanisms by which CDK1-cyclin A modifies IACs and the cytoskeleton.

The observation that CDK1-dependent phosphorylation of talin at S1589 alters the mechanical sensitivity of R7R8 has implications for the role of talin as a mechanosensitive signalling hub (27). This paradigm is based on the concept that the talin scaffold that connects integrins to the cytoskeleton can adopt many different conformations as a result of its binary switch patterns. These binary switches enable the cell to alter its behaviour in response to signalling, but also based on prior events, imbuing the cell with mechanical memory (51). The highly reproducible response of talin to mechanical force is due to each of the talin switch domains having different mechanical responses to force (44), and recruiting different signalling molecules as a function of force. The phosphorylation of S1589 by CDK1 may therefore alter the order that the talin switch domains unfold (Fig. 7). As further speculation, the concept of mechanical computation of the cell (51), in which the entire cytoskeletal architecture functions as a computational assembly, predicts that cell architecture is dictated by the signals that the cell has received and the current patterns of mechanical switches in the cells. Alteration of this pattern by talin S1589 phosphorylation might lead to global changes to mechanotransduction downstream of talin. It may be that talin signalling, mechanotransduction via mechanical linkages, and information storage in binary switches are controlled by PTMs across many different tissues and scenarios. In this context, cell polarity is a key driver of tissue self-organisation and coupling cell division to the geometry and mechanics of the cytoskeletal machinery may provide a means of regulating the polarity and organisation of cells into tissues that underpins multicellularity. Prior to division, most cells are attached to an underlying ECM via IACs, and/or to neighbouring cells. While modulation of adhesion occurs during interphase to enable mitotic cell rounding and physical separation of daughter cells, the links between the cell cycle and cell adhesion are poorly understood. The novel interaction between talin and CDK1 identified in this report provides a framework for understanding these linkages.

Our studies demonstrate that CDK1 is able to phosphorylate talin and switch talin into a state with different mechanical properties, potentially altering the talin interactome and function in IACs. As talin is central to the downstream events emanating from integrins, its interaction with CDK1 may enable large-scale, synchronised changes to cell architecture to be enabled during cell cycle progression and other adhesion-related processes such as cell migration. Furthermore, it is conceivable that changes in ECM stiffness sensed by talin may impact upon CDK1 activity and function, providing a direct, primordial link between mechanosensing and the induction of cell proliferation. Defining the changes in the talin and CDK1 interactomes that result from CDK1-talin binding, and the knock-on consequences for adhesion and cell cycle signalling will be a priority.

## Materials and methods

### Plasmid preparation and protein expression

DNA fragments encoding the talin R7R8 domain (residues 1361-1654 of TLN1_MOUSE), talin R7 (residues 1361-1462 and 1578-1658 of TLN1_MOUSE), and talin R7R8^S1589D^ (residues 1361-1654 of TLN1_MOUSE with an S1589D mutation) were synthesised by IDT block or PCR from template DNA. Each target DNA fragment was then assembled with a template vector (pET151: AviTag-(I27)_2_-Target-(I27)_2_-SpyTag) containing an AviTag at the N-terminus, a SpyTag at the C-terminus and four repeats of titin I27 domains as a molecular handle using NEBuilder HiFi DNA assembly. The sequence of all plasmids was verified by 1^st^ base sequencing service. Each plasmid was co-transformed with a BirA plasmid and expressed in *E. coli* BL21(DE3) cultured in LB-media with D-biotin, and affinity purified using the 6His-Tag.

### Antibodies and reagents

Monoclonal antibodies used were mouse anti-paxillin (clone 349; 1:500 for immunofluorescence; BD Biosciences; 610051), mouse anti-cyclin B1 (Clone GNS3; 1:2000; Merck Millipore 05-373), mouse anti-cyclin A2 (clone BF683; 1:1000, Cell Signaling Technology 4656), mouse anti-CDK1 (clone POH1; 1:1000; Cell Signaling Technology 9116), mouse anti-actin (clone AC-40; 1:2000; Sigma-Aldrich A3853), mouse anti-HA (clone 12CA5; 1:2000: Thermo Fisher MA1-12429), and rabbit anti-CDK1 pY15 (clone 10A11; 1:1000; Cell Signaling Technology 4539). Polyclonal antibodies used were rabbit anti-CDK1 (1:1000; Merck Millipore; ABE1403) and rabbit anti-CDK1 pT161 (1:1000; Cell Signaling Technology; 9114). Secondary Alexa-Fluor 680-conjugated (1:10,000; Thermo Fisher A10043) or DyLight 800-conjugated (1:10,000; Cell Signaling Technology 5257) antibodies were used for immunoblotting. Anti-mouse and anti-rabbit Alexa-Fluor 680-conjugated light chain-specific secondary antibodies were used (1:5000) for immunoblotting immunoprecipitations (Jackson ImmunoResearch; 115-625-174 and 211-622-171). Anti-mouse and anti-rabbit Alexa-Fluor 488- and 594-conjugated secondary antibodies (1:500) were used for immunofluorescence (all from Thermo Fisher). Thymidine and RO-3306 were purchased from Sigma-Aldrich. The cdc2-HA plasmid was obtained from Addgene (#188818).

### Protein purification

Expression constructs of talin were prepared using mouse talin-1 DNA cloned into the expression vector pET151-TOPO, expressed in *E.coli* BL21(DE3) Star and cultured in either minimal media for NMR samples or LB for non-labelled samples. Proteins were purified using nickel affinity chromatography and the His-tag removed by TEV protease cleavage (52). The protein was then further purified using ion exchange chromatography. The protein constructs used were mouse talin-1 R1-R3 (residues 482-911), R4-R8 (residues 913-1653), R7R8 (residues 1357-1653), R8 (residues 1461-1580), R9-R10 (residues 1655-1973), R10 (residues 1815-1973), R11-R12 (residues 1974-2294), R13-DD (residues 2300-254), and mouse talin-2 R7R8 (residues 1360-1656).

### Fluorescence polarisation

The following CDK1 peptides with a terminal cysteine residue were synthesised by GLBiochem (Shanghai): CDK1(206-223) GDSEIDQLFRIFRALGTP-C and CDK1-2A(206-223) GDSEAAQLFRIFRALGTP-C. BODIPY-TMR coupled peptides were dissolved and stored in PBS (137 mM NaCl, 27 mM KCl, 100 mM Na_2_HPO_4_, 18 mM KH_2_PO_4_, pH 7.4), 5 mM TCEP and 0.05% (v/v) Triton X-100. Uncoupled dye was removed using a PD-10 gel filtration column (GE Healthcare). Fluorescence polarisation measurements were recorded on a BMGLabTech CLARIOstar plate reader and analysed using GraphPad Prism (version 6.07). Kd values were calculated by nonlinear curve fitting using a one site total and non-specific binding model.

### NMR spectroscopy

NMR samples were prepared in 50 mM NaCl, 15 mM NaH_2_PO_4_, 6 mM Na_2_HPO_4_, 2 mM DTT, pH 6.5, 5% (v/v) D_2_O. All data were collected at a temperature of 298 K on a Bruker Avance III 600 MHz NMR spectrometer equipped with CryoProbe. The spectra were processed using Topspin (Bruker) and analysed using CCPN Analysis (53). Backbone resonance assignments of talin-1 R8 residues 1461-1580 were assigned previously (BMRB ID:19339 (33)).

### X-ray crystallography

Talin-1 R7R8 was concentrated to 300 μM in 50 mM NaCl, 3 mM DTT, 20 mM Tris, pH 7.4, and incubated with CDK1 peptide in an 8:1 molar excess. Sitting-drop sparse matrix crystallisation screening was performed using a Mosquito solution handling robot (TTP Labtech) with 400 nl drops and a 1:1 precipitate:precipitant ratio. Crystals formed in 20% Isopropanol, 20% PEG4K, 0.1 M Na citrate, pH 5.6 at 4°C in 3-4 weeks. Crystals were vitrified in mother liquor containing 25% (v/v) glycerol in liquid nitrogen. Diffraction data were collected on I24 (Diamond) and indexed and integrated using the Xia2 3dii pipeline in space group P212121. The structure was solved using molecular replacement using PHASER (54) with the search model 2X0C and two copies of R7R8 present in the asymmetric unit. Post molecular replacement electron density for the CDK1 peptide was visible in the F_0_-F_C_ difference map, allowing the unambiguous assignment of all side chain positions. The structure was refined using Phenix 1.17 (55) and modelled using COOT (56). Data were refined using isotropic B-factors, and prior to deposition with a round of weight optimisation of both the stereochemical and B-factor weights. Data reduction and refinement statistics are shown in Table 1 and the atomic coordinates were deposited to the PDB with accession code 6TWN.

### Cell culture, synchronisation and transfection

U2OS cells (European Collection of Cell Cultures 92022711; Sigma-Aldrich) were maintained in DMEM (Sigma-Aldrich) supplemented with 10% (v/v) FCS (Lonza), 1% (v/v) penicillin/streptomycin, and 2 mM l-glutamine at 37°C, 5% (v/v) CO_2_. For steady-state analysis of adhesion complexes in asynchronous cells, cells were cultured on glass coverslips for 48 h and then treated with indicated compounds for 1 h. U2OS cells were synchronized by using a double-thymidine block protocol. Cells were plated and after 24 h of growth, thymidine was added to a final concentration of 2 mM, and the cells were incubated for 16 h. Cells were then washed twice with PBS and allowed to grow for 8 h in fresh DMEM. Thymidine was then added to a final concentration of 2 mM for an additional 16 h before cells were washed twice with PBS and released into DMEM. U2OS cells were transfected with DNA constructs by using Lipofectamine 3000 reagent (Sigma-Aldrich) and siRNAs by using oligofectamine (Sigma-Aldrich) according to the manufacturer’s instructions. Knockdowns of CDK1 was performed by using SMARTpool reagents (L-003224-00-0005; GE Healthcare) and ON-TARGETplus nontargeting siRNA (GE Healthcare) was used as a negative control.

### GFP-Trap and mass spectrometry analysis

U2OS cells stably expressing talin-1-GFP or GFP were first generated via transient transfection using Lipofectamine 2000 (ThermoFisher Scientific) and selected using regular media supplemented with 500 μg/ml of G418. Cells were then plated on fibronectin for 2 hours, washed with PBS and lysed in CSK buffer [0.5% (w/v) Triton X-100, 10 mM Pipes, pH 6.8, 150 mM NaCl, 150 mM sucrose, 3 mM MgCl_2_, 10 mg/ml leupeptin, 10 mg/ml aprotinin, 0.5 mM 4-(2-aminoethyl) benzenesulfonyl fluoride hydrochloride, 2 mM Na_3_VO_4_]. Lysates were incubated with GFP-Trap agarose beads (Chromotek) for 1 hour at 4 ̊ C. Complexes bound to the beads were washed three times with ice-cold lysis buffer and eluted in Laemmli reducing sample buffer. Protein samples were separated by SDS-PAGE and following staining with InstantBlue (Expedeon), gel lanes were sliced into ten 2-mm bands and subjected to in-gel trypsin digestion (Humphries et al., 2009). Samples were analysed by LC-MS/MS using an UltiMate® 3000 Rapid Separation LC (RSLC, Dionex Corporation, Sunnyvale, CA) coupled to an Orbitrap Elite (Thermo Fisher Scientific, Waltham, MA) mass spectrometer. Peptide mixtures were separated using a gradient from 92% A (0.1% FA in water) and 8% B (0.1% FA in acetonitrile) to 33% B, in 44 min at 300 nl.min-1, using a 75 mm x 250 μm i.d. 1.7 MBEH C18, analytical column (Waters). Peptides were automatically selected for fragmentation by data-dependant analysis. Tandem mass spectra were extracted using extract_msn (Thermo Fisher Scientific) executed in Mascot Daemon (Matrix Science). Peak list files were searched against the SwissProt human database (version 3.70, May 2013). Carbamidomethylation of cysteine was set as a fixed modification, and oxidation of methionine was allowed as a variable modification. Only tryptic peptides were considered, with up to one missed cleavage permitted. Monoisotopic precursor mass values were used, and only doubly and triply charged precursor ions were considered. Mass tolerance for precursor and fragment ions was 0.5 Da. Data were validated in Scaffold using the following threshold: at least 80% probability at the peptide level, at least 99% probability at the protein level and at least two unique peptides. Relative protein abundance was calculated using the unweighted spectral count of a given protein normalised to the total number of spectra observed in the entire sample and to the molecular weight of that protein (normalised spectral count). SAINT and FC_B scores were calculated using the Resource for Evaluation of Protein Interaction Networks (REPRINT) online tool (https://reprint-apms.org/). The mass spectrometry proteomics data have been deposited to the ProteomeXchange Consortium via the PRIDE (57) partner repository with the dataset identifier PXD024634 and 10.6019/PXD024634.

### Proximity biotinylation, purification of biotinylated proteins and mass spectrometry analysis

U2OS cells were seeded onto plastic tissue culture plates overnight, then transfected with BirA empty vector or BirA-CDK1 and incubated for a further 16 hours. Transfected cells were then incubated in medium with 50 μM biotin for 24 h. Biotinylated proteins were affinity purified following a protocol adapted from Roux *et al*. (58, 59). Three 10-cm plates of cells were washed three times in PBS, and cells were lysed with 400 μl lysis buffer (50 mM Tris-HCl, pH 7.4, 250 mM NaCl, 0.1% [w/v] SDS, 0.5 mM DTT, and 1× cOmplete Protease Inhibitor Cocktail) at RT. 120 μl 20% (v/v) Triton X-100 was added, and cell lysates were maintained at 4°C. DNA was sheared by passing cell lysates through a 19G needle four times before 360 μl chilled 50 mM Tris-HCl, pH 7.4, was added, and then passing through a 27G needle four times. Cell lysates were centrifuged at full speed for 10 min at 4°C, and supernatant was rotated with 45 μl MagReSyn streptavidin beads (ReSyn Biosciences) at 4°C overnight. Beads were washed twice with 500 μl wash buffer 1 (10% [w/v] SDS), once with 500 μl wash buffer 2 (0.1% [w/v] deoxycholic acid, 1% [w/v] Triton X-100, 1 mM EDTA, 500 mM NaCl, and 50 mM HEPES), and once with 500 μl wash buffer 3 (0.5% [w/v] deoxycholic acid, 0.5% [w/v] NP-40, 1 mM EDTA, and 10 mM Tris/HCl, pH 7.4). Proteins were eluted in 40 μl of 2× reducing sample buffer with 100 μM biotin for 10 min at 70°C. Eluted proteins were briefly subjected to SDS-PAGE (3 min at 200 V, 4– 12% Bis-Tris gel, Life Technologies) and stained with InstantBlue Coomassie protein stain before being washed with ddH_2_O overnight at 4°C. Bands were excised and transferred to wells in a perforated 96-well plate, and in-gel tryptic digestion was performed as previously described (60). Peptides were desalted using 1 mg POROS Oligo R3 beads (Thermo Fisher Scientific). Beads were washed with 50 μl 0.1% (v/v) formic acid (FA) before the peptide solution was added. Beads were washed twice with 100 μl 0.1% (v/v) FA, and peptides were eluted twice with 50 μl 50% (v/v) acetonitrile (ACN) and 0.1% (v/v) FA. Peptides were dried using a vacuum centrifuge and resuspended in 11 μl 5% (v/v) ACN and 0.1% (v/v) FA. Peptides were analysed by liquid chromatography (LC)–tandem MS (MS/MS) using an UltiMate 3000 Rapid Separation LC (RSLC, Dionex Corporation) coupled to an Orbitrap Elite mass spectrometer (Thermo Fisher). Peptides were separated on a bridged ethyl hybrid C18 analytical column (250 mm × 75 μm inner diameter, 1.7 μm particle size, Waters) over a 1 h gradient from 8 to 33% (v/v) ACN in 0.1% (v/v) FA. LC–MS/MS analyses were operated in data-dependent mode to automatically select peptides for fragmentation by collision-induced dissociation (CID). Quantification was performed using Progenesis LC–MS software (Progenesis QI, Nonlinear Dynamics; http://www.nonlinear.com/progenesis/qi-for-proteomics/). In brief, automatic alignment was used, and the resulting aggregate spectrum filtered to include +1, +2 and +3 charge states only. A.mgf file representing the aggregate spectrum was exported and searched using Mascot (one missed cleavage, fixed modification: carbamidomethyl [C]; variable modifications: biotinylation [B], oxidation [M]; peptide tolerance: ± 5 ppm; MS/MS tolerance: ±0.5 Da), and the resulting .xml file was re-imported to assign peptides to features. Three separate experiments were performed, and abundance values for proteins identified in the analysis were used to determine which proteins were enriched over twofold in cells expressing BirA-CDK1 relative to cells expressing BirA-empty vector control. The mass spectrometry proteomics data have been deposited to the ProteomeXchange Consortium via the PRIDE (57) partner repository with the dataset identifier PXD024796 and 10.6019/PXD024796.

### Immunoprecipitation

U2OS cells transfected with GFP-tagged talin-1, GFP-tagged ΔR8 talin-1 or GFP alone or synchronized in G2 9 hours after thymidine arrest (two 15-cm-diameter dishes per condition) were lysed (500 μl per dish) in modified CSK buffer (150 mM NaCl, 25 mM Tris-HCl, pH 7.4, 1 mM EDTA, 1% (v/v) NP-40), protease inhibitors (11836145001; Roche) and PhosStop reagent (Roche). Lysates were passed five times through a narrow-bore tip before centrifugation (10000 *g* for 3 min at 4°C). After centrifugation, 20 μl GFP-trap Sepharose beads (Chromotek) or immunoprecipitating mAbs (mouse anti–cyclin B1 clone GNS3 or mouse anti–cyclin A clone E67.1; SC53230; Santa Cruz Biotechnology, Inc., or mouse IgG; Sigma-Aldrich) were added to the lysate (2 μg/ml cyclin B1 and 10 μg/ml cyclin A final concentration) together with protein G Sepharose (20 μl of 50% slurry bead volume; GE Healthcare) for 16 h at 4°C. Sepharose beads were then collected and washed two times in lysis buffer and once in distilled H_2_O by centrifugation (2,800 *g* for 2 min). GFP-associated or immunoprecipitated complexes were eluted (30 μl) at pH 2 for 5 min at 25°C and neutralized according to the manufacturer’s instructions (88805; Thermo Fisher Scientific). Samples were then reduced at 70°C for 5 min by dilution in 5x sample buffer (125 mM Tris, 10% (w/v) SDS, 25% (v/v) glycerol, 0.01% (w/v) bromophenol blue, and 10% (v/v) β-mercaptoethanol) and subjected to SDS-PAGE and Western blotting by using the Odyssey Infrared Imaging System. To avoid detection of antibody heavy chains, light chain–specific Alexa Fluor 680–conjugated secondary antibodies were used (1:5,000).

### Immunofluorescence microscopy

U2OS cells were fixed in 4% (w/v) PFA for 20 min, washed twice with PBS, and permeabilized by using 0.2% (w/v) Triton X-100 in PBS for 10 min. Cells were then washed with PBS and PFA quenched by incubation with 0.1 M glycine/PBS for 30 min. Cell were washed with PBS three times and then incubated with primary antibodies (45 min at RT) and washed with PBS containing 0.1% (w/v) Tween-20 (PBST) and incubated for 30 min with the appropriate secondary antibodies and, where applicable, Alexa Fluor dye–conjugated phalloidin (Thermo Fisher Scientific). Finally, cells were washed three times with PBST and once with distilled H_2_O before being mounting on coverslips by using ProLong diamond antifade reagent (Thermo Fisher Scientific) and imaging. Images were acquired on an inverted confocal microscope (TCS SP5 Acousto-Optical Beam Splitter; Leica Microsystems) by using a 63x objective (HCX Plan Apochromat, NA 1.25) and Leica Confocal Software (Leica Microsystems), and image analysis was performed using ImageJ. Images were background subtracted by using rolling ball subtraction, and images of paxillin staining (mouse anti-paxillin, clone 349, BD Biosciences) or GFP-talin-1 were thresholded to define adhesion complexes. By using a size cut-off of 0.2 μm, the total area of adhesion complexes was determined per cell as a proportion of total cell area. Representative cells were selected based on consistency with phenotype observed across the field of view, with distinct cells present within a similar density of surrounding cells being chosen for analysis.

### Total Internal Reflection Fluorescence (TIRF) microscopy

U2OS cells stably expressing GFP-talin-1 constructs were transfected with mScarlet-CDK1 using Lipofectamine 3000 according to manufacturer’s instructions. Cells were then plated onto glass-bottomed dishes (Mat-tek), cultured overnight, and fixed with 4% (w/v) PFA for 20 min then washed twice with PBS. Images were collected on a Leica Infinity TIRF microscope using a 100x/1.47 HC PL Apo Corr TIRF Oil objective with 488 nm and 561 nm diode TIRF lasers (with 30% laser power and a penetration depth of 110 nm) and an ORCA Flash V4 CMOS camera (Hamamatsu) with 300 ms exposure time and camera gain of 2. Images were background subtracted by using rolling ball subtraction using ImageJ and representative images were selected based on consistency with phenotype observed across the field of view.

### Immunoblotting

Cells were lysed in lysis buffer (150 mM NaCl, 25 mM Tris-HCl, pH 7.4, 1 mM EDTA, 1% (v/v) NP-40, 5% (v/v) glycerol, 1x protease inhibitor cocktail [Sigma Aldrich], and 1× PhosSTOP phosphatase inhibitor cocktail [Sigma-Aldrich]). Lysates were clarified by centrifugation at 10,000 *g* for 5 min at 4°C. Cell lysates were separated by SDS-PAGE (4–12% Bis-Tris gels; Thermo Fisher Scientific) under reducing conditions and transferred to nitrocellulose membranes (Whatman). Membranes were blocked for 60 min at RT using 5% (w/v) BSA in TBS (10 mM Tris-HCl, pH 7.4, and 150 mM NaCl) containing 0.05% (w/v) Tween-20 (TBST) and then probed overnight with primary antibodies diluted in 5% (w/v) BSA/TBST at 4°C. Membranes were washed for 30 min by using TBST and then incubated with the appropriate fluorophore-conjugated secondary antibody diluted in 5% (w/v) BSA/TBST for 45 min at RT in the dark. Membranes were washed for 30 min in the dark by using TBST and then scanned by using the Odyssey infrared imaging system (LI-COR Biosciences)

### In vitro kinase assay

Purified recombinant GST-tagged CDK1-cyclin A2 and His_6_-tagged CDK1-cyclin B1 (Invitrogen) were stored in 150 mM NaCl, 0.5 mM EDTA, 0.01% (v/v) Triton X-100, 2 mM DTT, 20% (v/v) glycerol 20 mM Tris, pH 7.5. 30 ng of each protein was mixed with 0.1 μg of substrate (talin R7R8) and incubated in 150 mM NaCl, 5 mM EDTA, 5 mM DTT, 25 mM MgCl_2_, 0.02% (v/v) Triton X-100, 1 mM ATP, 50 mM HEPES, pH 7.4 at 30 °C with shaking at 100 rpm for 20 min. The reaction was stopped by adding SDS sample buffer and boiled at 95 °C for 10 min. A gradient SDS-PAGE 4-12% Bis-Tris gel (Thermo Fisher Scientific) was loaded with the entire sample and run at 200 V for 45 min. in 1x SDS running buffer (NuPAGE). After western blotting, anti-phosphorylated CDK substrate antibody (Cell Signalling Technology) was used to probe the reactions.

### Phosphorylation site analysis

Phosphorylation sites of talin-1 R7R8 and talin-2 R7R8 were determined using mass spectrometry. The *in vitro* kinase assay was carried out with the substrates talin-1 R7R8, talin-2 R7R8 and CDK1-cyclinA2. Reactions was carried out for 45 min. at 30 °C and stopped by adding SDS sample buffer and boiling for 10 min at 95 °C. All samples were loaded onto an SDS PAGE 4-12% Bis-Tris gel (Thermo Fisher Scientific) and separated by running at 200 V for 60 min. Gels were stained with Instant-Blue (Expedeon) for 15 min, and washed in water overnight at 4°C. The talin R7R8 bands were cut from the gel and processed by in-gel tryptic digestion. Peptides were analysed by LC-MS/MS by using an UltiMate 3000 Rapid Separation LC (Dionex Corporation) coupled to an Orbitrap Elite MS (Thermo Fisher Scientific). Peptides were separated on a bridged ethyl hybrid C18 analytical column (250 mm × 75 μm internal diameter, 1.7 μm particle size; Waters) over a 45 min gradient from 8 to 33% (v/v) acetonitrile in 0.1% (v/v) formic acid. LC-MS/MS analyses were operated in data-dependent mode to automatically select peptides for fragmentation by collision-induced dissociation. Multistage activation was enabled to fragment product ions resulting from neutral loss of phosphoric acid. Quantification was performed using Progenesis LC-MS/MS software.

### Single molecule stretching

All single molecule stretching experiments were performed using magnetic tweezers (61–63) in standard buffered solution comprising PBS, 1% (w/v) bovine serum albumin, 2 mM dithiothreitol, 10 mM sodium L-ascorbate at 21±1 °C. The height of the target molecule-tethered superparamagnetic bead from the coverslip surface was recorded. At a constant applied force, the bead height change was the same as the molecule extension change (64). During linear force-increase/force-decrease scans with typical loading rates of 0.1 to 10 pN s^−1^, the stepwise bead height change was the same as the stepwise extension change of the molecule. Over the time window of the stepwise transition event (≤0.01 s, the temporal resolution of this setup), the force change (≤0.001 to 0.1 pN) was negligible. Force calibration of the magnetic tweezer has ~10% uncertainty due to the heterogeneity of the paramagnetic beads (61).

Each single molecule protein construct contained a biotinylated AviTag at its N-terminus, a target domain spanned between four repeats of titin I27 domains, and a SpyTag at its C-terminus (65). Between each two neighbouring molecular components in the construct, a short linker, GGGSG, was included to ensure flexibility of the components. The C-terminus of the protein construct was specifically attached to a SpyCatcher-coated coverslip surface (Paul Marienfield), and the N-terminus of the construct was attached to a biotinylated DNA-coated paramagnetic bead (2.8 μm diameter, Invitrogen) via a biotin-neutravidin interaction. The four repeats of I27 domains and the DNA handle acted as a molecular spacer to avoid non-specific bead-surface interactions. The I27 domain has an ultra-high mechanical stability in that it typically unfolds at >100 pN with a characteristic step size of ~24 nm at force loading rates 1-5 pN s^−1^ (66). Hence it can be distinguished from the target domain unfolding signals and does not affect the probing and characterisation of the target domains.

## Acknowledgements

We thank David Critchley for critical reading of the manuscript.

## Supplementary data

***Supp Table 1***

GFP-Trap data for GFP-talin-1 – See accompanying spreadsheet

***Supp Table 2***

BioID for BirA-CDK1– See accompanying spreadsheet

**Supp Table 3.** Data collection and refinement statistics for Talin1 R7R8 in complex with CDK1(206-223)

|  |  |
| --- | --- |
| *Beamline* | I24 (Diamond) |
| Wavelength (Å) | 0.9686 |
| Resolution range (Å) | 46.15-2.28 (2.35-2.28) |
| Space group | P2<sub>1</sub>2<sub>1</sub>2<sub>1</sub> |
| Unit Cell |  |
| a,b,c (Å) | 69.00 97.87 104.67 |
| $\alpha,\beta,\gamma$ (°) | 90 90 90 |
| Unique reflections | 32821 |
| Completeness (%) | 99.6 (98.9) |
| Multiplicity | 5.1 (4.8) |
| CC1/2 | 0.992 (0.541) |
| I/$\sigma$(I) | 7.0 (1.2) |
| R<sub>merge</sub> (I+/I-) | 0.204 (1.309) |
| R<sub>pim</sub> (I+/I-) | 0.099 (0.652) |
| Refinement |  |
| R<sub>work</sub> (%) | 21.44 (30.17) |
| R<sub>free</sub> (%) | 26.18 (33.90) |
| No. of atoms |  |
| Macromolecule | 4813 |
| Solvent | 180 |
| RMSD bonds (Å) | 0.003 |
| RMSD angles (°) | 0.521 |
| Ramachandran favoured (%) | 99.38 |
| Ramachandran allowed (%) | 0.46 |

R-free was calculated using 5% of all reflections isolated from refinement. Data was deposited to the PDB with the accession code 6TWN.

**Supp Fig 1:**
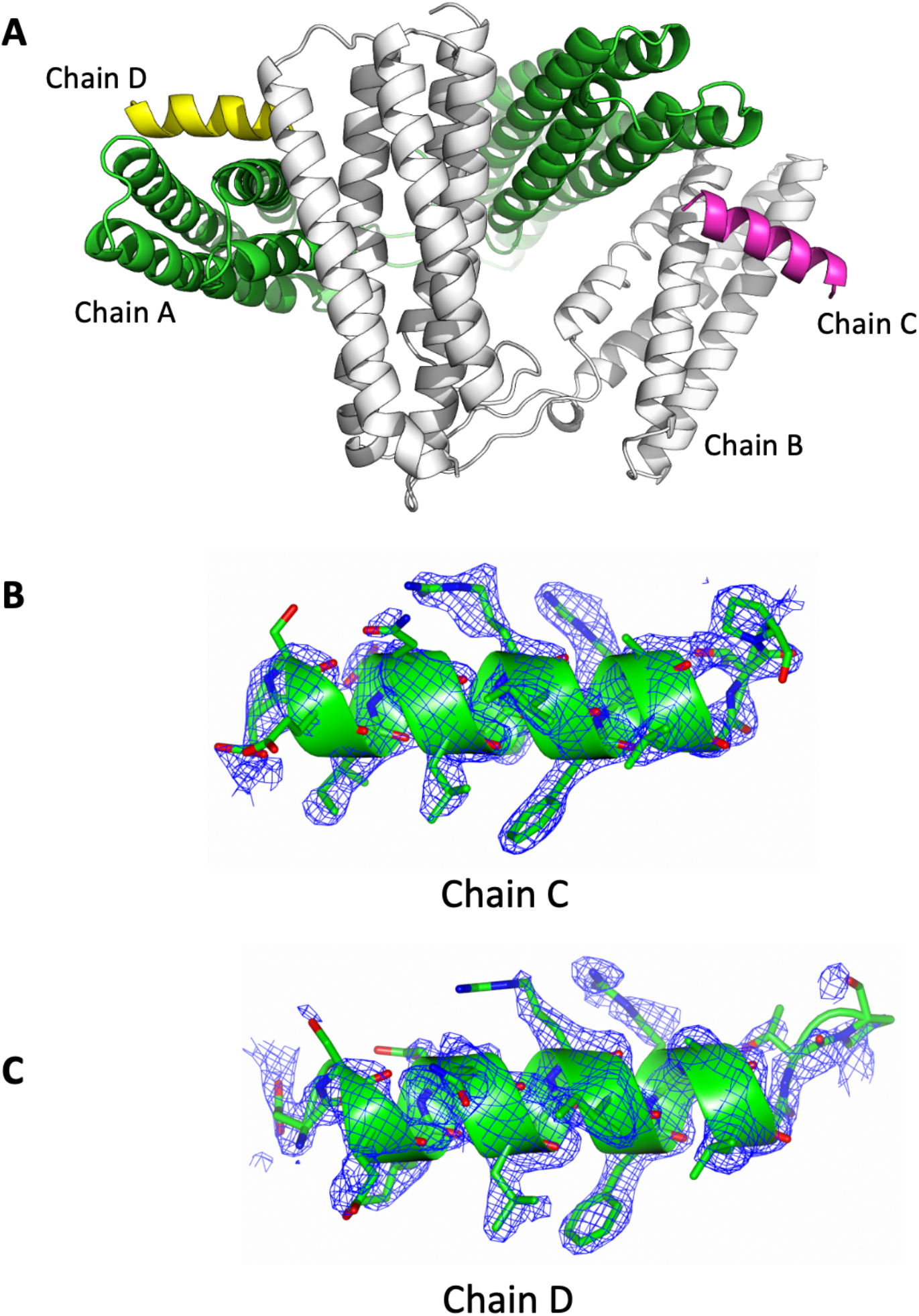
Structural characteristics of the R7R8-CDK1 complex. (**A**) Cartoon representation of the two R7R8-CDK1 complexes in the asymmetric unit. (**B-C**) Simulated annealing composite omit map of CDK1 peptides contoured of 1σ in blue. (**B**) Chain C and (**C**) Chain D.

**Supp Fig 2:**
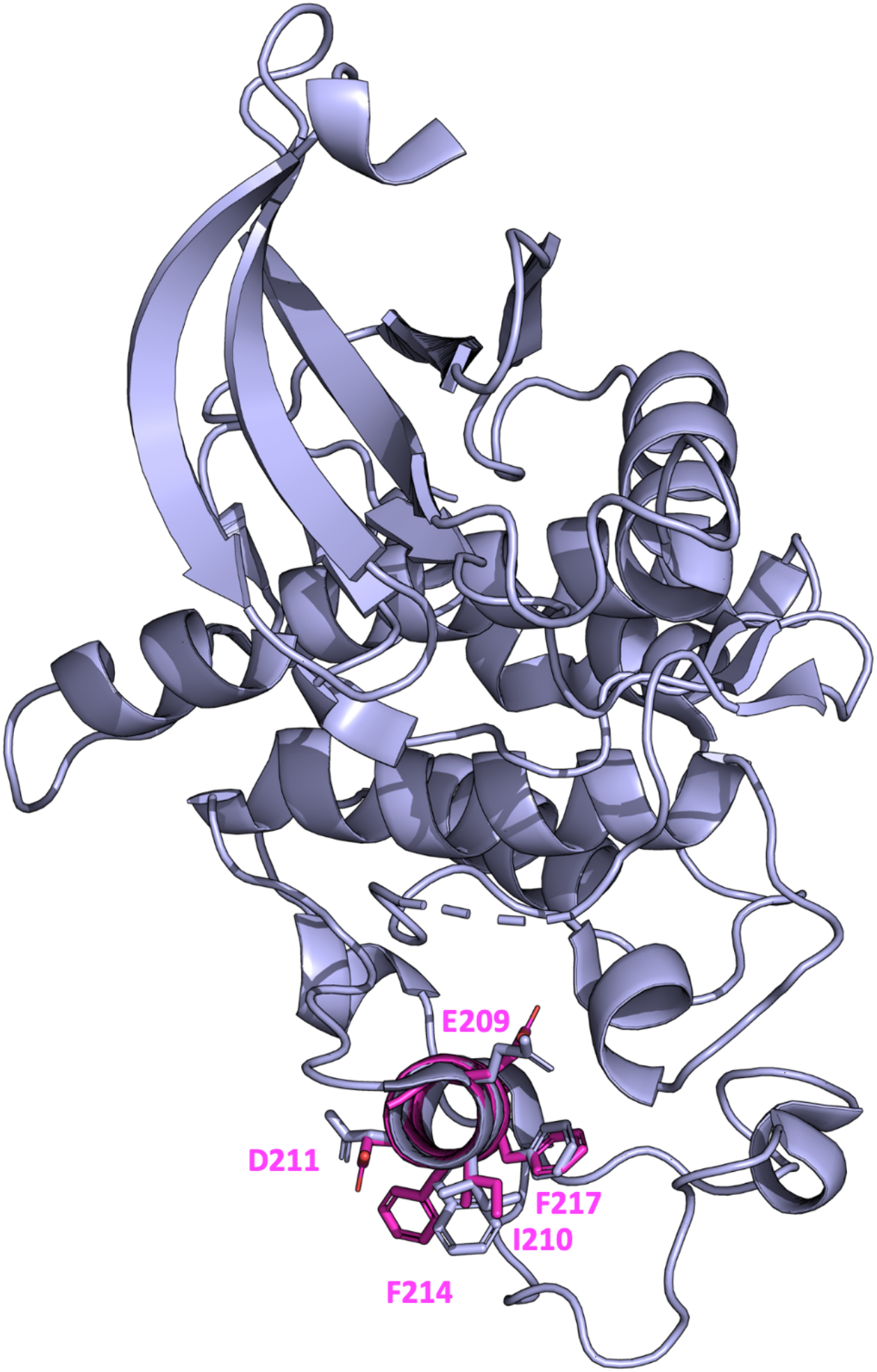
Overlay of CDK1 with the CDK1 LD motif in the talin-bound form. (**A**) CDK1 structure (blue) PDB ID: 4YC6 overlaid with CDK1 peptide (pink) from the talin:CDK1 complex PDB ID: 6TWN.

**Supp Fig 3:**
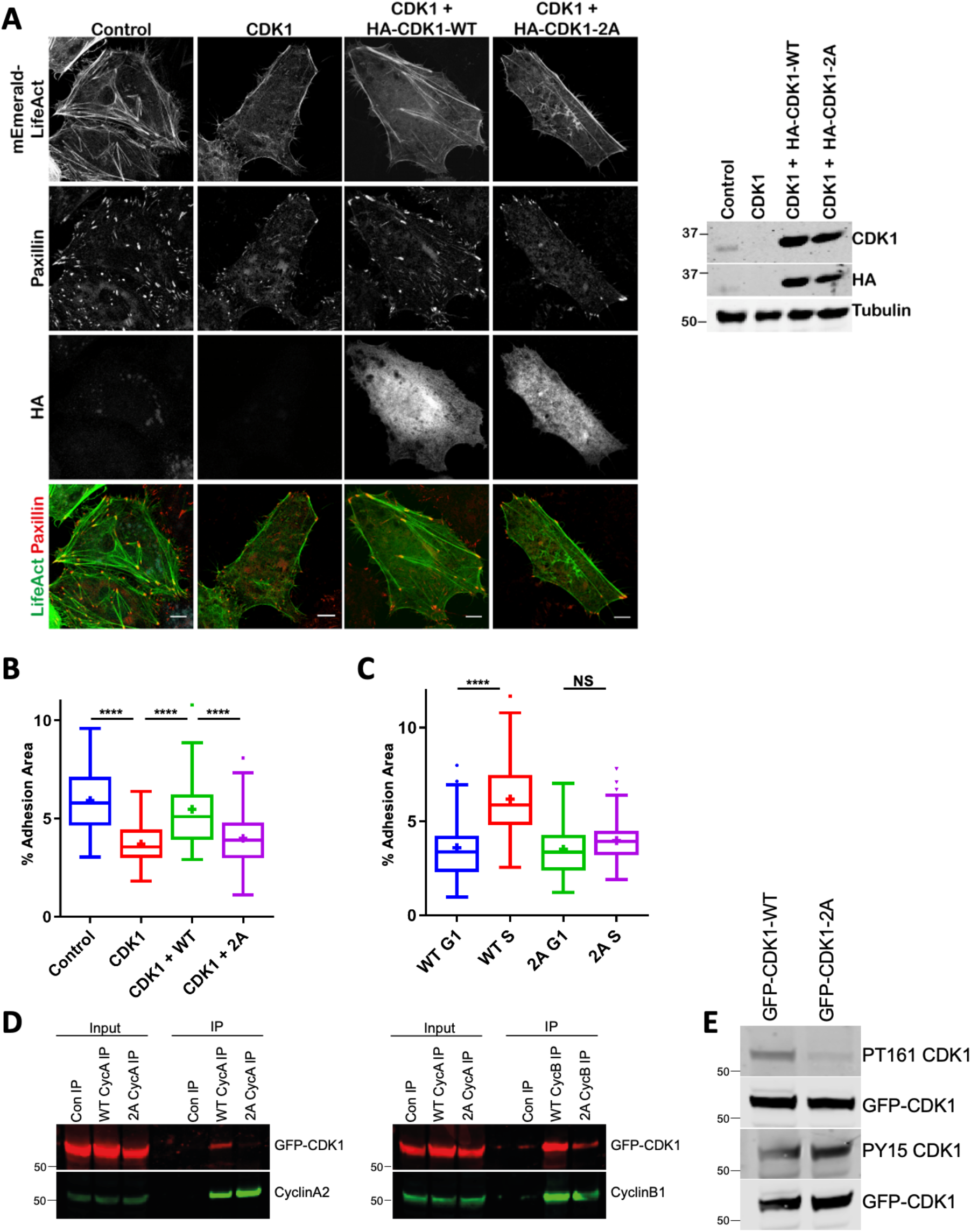
CDK1-2A mutant perturbs CDK1-dependent regulation of IACs and CDK1 binding to cyclin A2. (**A**) Immunofluorescence images of control, CDK1-knockdown cells, and CDK1-knockdown cells re-expressing WT CDK1 or CDK1-2A stained for paxillin, actin and HA. Bars 10 μm. (right) Western blot confirming the knock down of CDK1 and re-expression of HA-tagged CDK1-WT and CDK1-2A. (**B**) Quantification of IAC area per cell after CDK1 knockdown and re-expression. A minimum of 50 cells per condition was used for analysis. Bars, 10 μm. (**C**) Quantification of changes in IAC area per cell in G1, S, and G2 phase for cells expressing WT CDK1 or CDK1-2A. A minimum of 50 cells per condition was used for analysis. Results in B and C are displayed as Tukey box and whisker plots (whiskers represent 1.5x interquartile range). ****, p<0.0001. (**D**) Immunoprecipitation of cyclin A2 (left) or cyclin B1 (right) from G2-synchronised cells expressing GFP-tagged CDK-WT or CDK1-2A and western blotted for GFP or cyclin. (**E**) Immunoprecipitation of GFP-CDK1-WT and GFP-CDK1-2A expressed in cells followed by western blotting for phosphorylation of T161 and Y15.

**Supp Fig 4:**
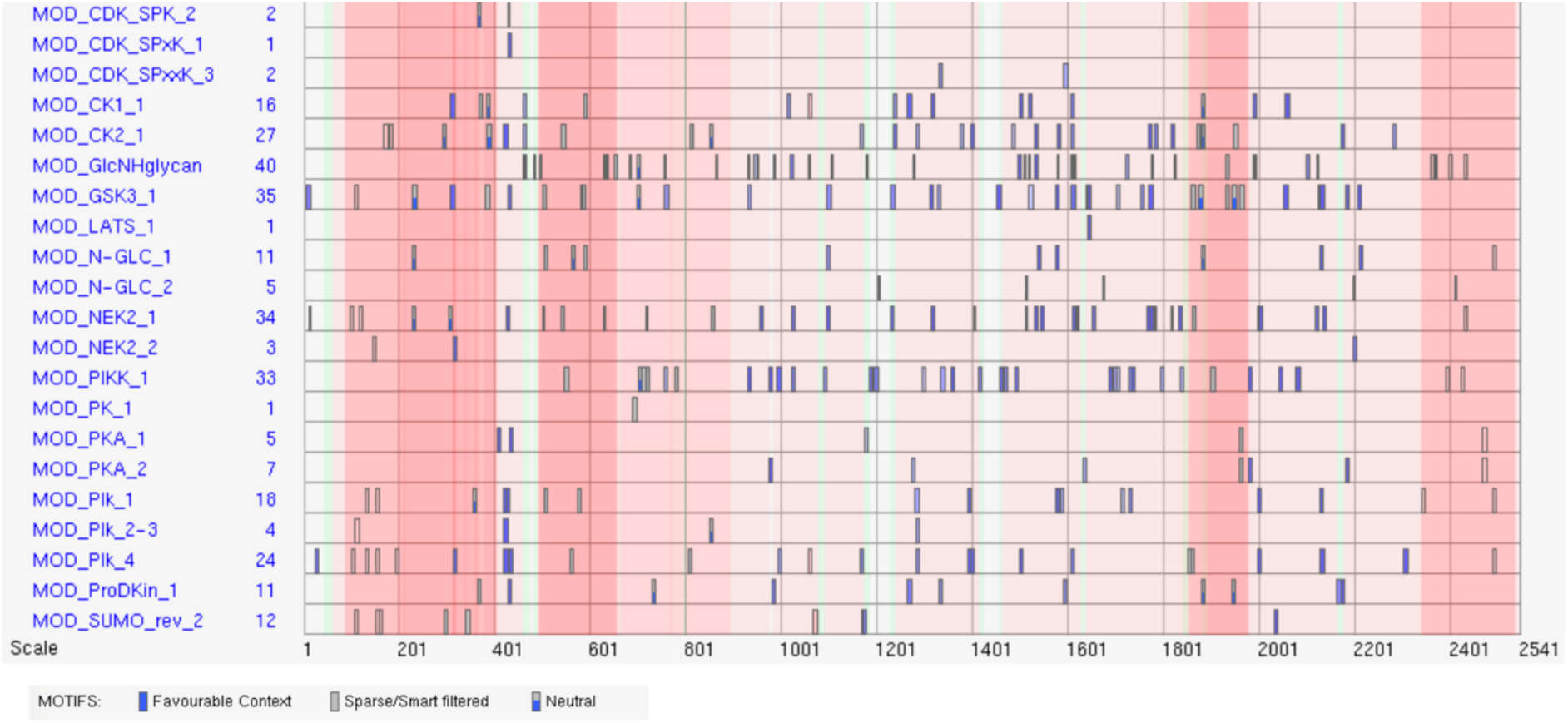
The posttranslational modifications of talin as indexed in the ELM database. The Eukaryotic Linear Motif (ELM) resource for functional sites in proteins (67) predicts that the talin switch domains are heavily modified by posttranslational modifications. The table shows part of the ELM analysis for mouse talin-1 (UniProt ID P26039). The rectangles denote the identified short linear motifs that fit the consensus linear motifs for different enzymes.

